# Evolutionary Co-Option of an Ancestral Cloacal Regulatory Landscape During the Emergence of Digits and Genitals

**DOI:** 10.1101/2024.03.24.586442

**Authors:** Aurélie Hintermann, Christopher Chase Bolt, M. Brent Hawkins, Guillaume Valentin, Lucille Lopez-Delisle, Sandra Gitto, Paula Barrera Gómez, Bénédicte Mascrez, Thomas A. Mansour, Tetsuya Nakamura, Matthew P. Harris, Neil H. Shubin, Denis Duboule

## Abstract

The transition from fins to limbs has been a rich source of discussion for more than a century. One open and important issue is understanding how the mechanisms that pattern digits arose during vertebrate evolution. In this context, the analysis of *Hox* gene expression and functions to infer evolutionary scenarios has been a productive approach to explain the changes in organ formation, particularly in limbs. In tetrapods, the transcription of *Hoxd* genes in developing digits depends on a well-characterized set of enhancers forming a large regulatory landscape^1,2^. This control system has a syntenic counterpart in zebrafish, even though they lack *bona fide* digits, suggestive of deep homology^3^ between distal fin and limb developmental mechanisms. We tested the global function of this landscape to assess ancestry and source of limb and fin variation. In contrast to results in mice, we show here that the deletion of the homologous control region in zebrafish has a limited effect on the transcription *of hoxd* genes during fin development. However, it fully abrogates *hoxd* expression within the developing cloaca, an ancestral structure related to the mammalian urogenital sinus. We show that similar to the limb, *Hoxd* gene function in the urogenital sinus of the mouse also depends on enhancers located in this same genomic domain. Thus, we conclude that the current regulation underlying *Hoxd* gene expression in distal limbs was co-opted in tetrapods from a preexisting cloacal program. The orthologous chromatin domain in fishes may illustrate a rudimentary or partial step in this evolutionary co-option.

## INTRODUCTION

The organization of tetrapod limbs has been conserved since their origin, with a universal pattern of ‘segments’ along the proximal to distal axis. The stylopod is a single bone (upper arm, leg) attached at one end to the torso and at the other end to two zeugopod bones (lower arm, leg), then, most distal are the mesopod (wrist, ankle) and the autopod (hand, foot). The formation of this generic pattern began before the water-to-land transition as sarcopterygian fishes display structures clearly related to proximal tetrapod limb structures^4^. However, when homologies are considered between fin structures and the most distal parts of tetrapod appendages, the mesopod and the autopod, there remains debate if fishes possess homologous skeletal rudiments. While mesopodial elements and extensive distal segments are present in sarcopterygian fins, the presence of true digital homologues has remained controversial^5,6^.

Because the *HoxA* and *HoxD* gene clusters were shown to be instrumental in making tetrapod limbs^7–10^, their expression domains during fin development were used to infer the presence of an autopod-related structure in fishes. In particular, *Hoxa13* and *Hoxd13* were studied due to their specific autopodial expression in tetrapod limbs^10^ and because their combined inactivation in mice produce autopodial agenesis^7^. An analysis of *hox13* genes in the distal teleost fin suggested that a ‘distal program’ also exists in fishes, implying that this genetic regulatory network, or part thereof, would have preceded digit formation in tetrapods^11,12^. This core and potentially ancestral distal pattern is however realized in formation of the dermal rays of fins, while the concurrent rudimentary endoskeleton, an array of singular radials connected to the girdle, was hypothesized to be primarily proximal^5^. In such a scenario, the autopods of tetrapods are proposed to form from the postaxial vestiges of an ancestral sarcopterygian fin^13,14^. The partial retention of expression patterns that presage the emergence of digits in ray-finned and chondrichthyan fishes is nevertheless suggestive of a common regulatory program shared amongst vertebrates, the deployment of which in different species accompanied changes in form^13^.

During tetrapod limb bud development, a series of enhancers within in a large regulatory landscape positioned 3’ of the *HoxD* gene cluster (3DOM) control the transcription of *Hoxd* genes up to *Hoxd11* in a proximal expression domain. These expression domains encompass tissue of the future stylopod (arm) and zeugopod (forearm) (Fig. S1a, green and schemes on the left)^15^. Posterior-distal limb bud cells then switch off these enhancers and activate another large regulatory landscape (5DOM), located 5’ to the gene cluster. This region is enriched with conserved enhancer elements that have been found to control the formation of digits by activating *Hoxd13* and its closest neighbors (Fig. S1a, blue). While the deletion of 3DOM abrogated the expression of all *Hoxd* genes in the proximal limb domain^15^, deletion of 5DOM removed all *Hoxd* mRNAs from the forming autopod^2^.

In the orthologous zebrafish *hoxda* cluster, genes are also expressed during early fin bud development, with progressively nested expression domains comparable to the murine situation^13,16^. At a later stage, transcription of both *hoxd9a* and *hoxd10a* persists in the ‘preaxial’ (anterior) part of the fin bud only (Fig. S1b, magenta), while *hoxd11a*, *hoxd12a* and *hoxd13a* transcripts are restricted to ‘postaxial’ (posterior) cells (Fig. S1b, orange), as is the case in the emerging fin bud^16^. For the latter genes, combined inactivation have revealed their function during distal fin skeletal development^11,17^. However, despite the analysis of distal enhancers orthologous to those of the mouse^12^, the functionality of the complete 3DOM and 5DOM regulatory landscapes in zebrafish has not been addressed. Hence, the existence of a comparable bimodal regulation of *Hoxd* genes has remained elusive, precluding any conclusion on its evolutionary origin.

We asked if the zebrafish *hoxda* genes were regulated by a comparable enhancer hub located at a distance from the gene cluster in a manner similar to mouse limbs^12,18^ (Fig. S1b, question marks). By deleting the orthologous zebrafish *hoxda* regulatory landscapes, we find that while the proximal appendage regulation by 3DOM is fully conserved between fish and mice, the core long-distance regulation by 5DOM underlying the distal expression is deficient in fins. The deletion of the zebrafish 5DOM revealed however that it shared with mouse an ancestral function in patterning the cloacal area. Our findings suggest this core ancestral regulatory landscape arose first in the ancient cloaca and was subsequently redeployed during the evolution and shaping of tetrapod digits and external genitals.

## RESULTS

### The zebrafish *hoxda* locus

The zebrafish *hoxda* locus shares a high degree of synteny with that of the *HoxD* locus in mammals, reflecting broad conservation given the key patterning role of this complex in development of many axial structures. The gene cluster is flanked by two gene deserts referred to as 3DOM (3’-located domain) and 5DOM (5’-located domain). As in mammals, the extents of both 3DOM and 5DOM correspond to topologically associating domains (TADs) and 3DOM is split into two sub-TADs (Fig. S2). This remarkable similarity in 3D conformations, though with a 2.6-fold difference in size between the mouse *versus* zebrafish locus, is further supported by the conserved position and orientation of critical CTCF binding sites within the gene clusters and their enrichment at TAD and sub-TAD borders (Fig. S2).

Interspecies genomic alignments reveal several conserved sequences within 5DOM across vertebrates, whereas little conservation was scored in 3DOM (Fig. S3a, b). Within the 5DOM comparison, we identified several of the previously annotated mouse enhancers in their zebrafish counterpart^12,19^. Consistent with the apparent conservation of chromatin structure, we found the same global organization of both coding and non-coding elements as in the mouse landscape. When compared to the size of the *Hox* cluster, the relative sizes of both gene deserts are bigger in mouse than in fish, and the zebrafish 5DOM was found to be larger than 3DOM, opposite to the mouse situation (Fig. S2c). Since the overall genomic organization of both *HoxD* loci is well conserved between mammals and fishes, we have concluded that these two flanking gene deserts and their topologically associating domains are ancestral features predating the divergence between ray finned fishes and tetrapods, likely conserved due to important regulatory functions. Whether these domains have, or retain *Hox* gene regulation as initially defined at this locus in the mouse^1,20^ remained nevertheless unclear.

### Regulatory potential of zebrafish *hoxda* flanking gene deserts

To address the potential function(s) of fish *hoxd* gene deserts, we explored chromatin accessibility and histone modification profiles using ATAC-seq^21^ and CUT&RUN^22^ assays, respectively, with posterior trunk as a source of cells, i.e., a domain where most *hox* genes are active. As a control sample, we used corresponding dissected heads where *hoxda* genes are not expressed (Fig. S4a, b). This analysis revealed enriched ATAC-seq signals not only within the *hoxda* cluster but also in the two gene deserts, with stronger signals in the 3DOM region (Fig. S4c). This suggested that in zebrafish, both gene deserts may indeed serve as regulatory landscapes, with long-range acting enhancers as is the case in tetrapods. Histone profiling supported this hypothesis with the same regions enriched in H3K27ac marks, while showing a poor (if any) enrichment in the negative H3K27me3 marks, suggesting several chromatin segments engaged in active transcriptional regulation.

To assess the functional potential of both *hoxda* gene deserts, we generated zebrafish mutant lines carrying full deletions either of 5DOM (*hoxda^de^*l(5DOM), referred to as Del(5DOM) below), or of 3DOM (*hoxda^del^*(3DOM) or Del(3DOM) below), using CRISPR-Cas9 chromosome editing. We first examined the impact of these large deletions on *hoxd13a, hoxd10a* and *hoxd4a* expression using whole-mount *in situ* hybridization (WISH), spanning from 30 hours post fertilization (hpf), i.e., from the onset of *hoxd13a* expression^16^ to 60 hpf and 72 hpf, stages when all *hoxda* gene expression decreases significantly in fin buds. In Del(3DOM) mutant embryos, expression of both *hoxd4a* and *hoxd10a* completely disappeared from the pectoral fin buds (Fig. 1a, right and middle panels, arrowheads). The same effect was observed at all stages analyzed (Fig. 1a). These data are consistent with similar analysis in mice where the limb proximal expression domain was no longer visible upon deletion of the 3DOM landscape^15^. This demonstrates that, as in tetrapods, enhancers controlling the transcription of *hoxd3a* to *hoxd10a* during fin bud development are located in the adjacent 3’ landscape. The 3DOM thus has an ancestral regulatory function in the development of proximal paired appendages. Expression of *hoxd13a* in post-axial cells, however, remained unchanged, with a global transcript distribution indistinguishable from wild-type fin buds (Fig. 1a, left panels, arrowheads). These data indicate that the control of *hoxd13a* expression is distinct from that impacting *hoxd3a* to *hoxd10a,* as is also the case in tetrapods^2^ (Fig. S1a).

**Figure 1.**
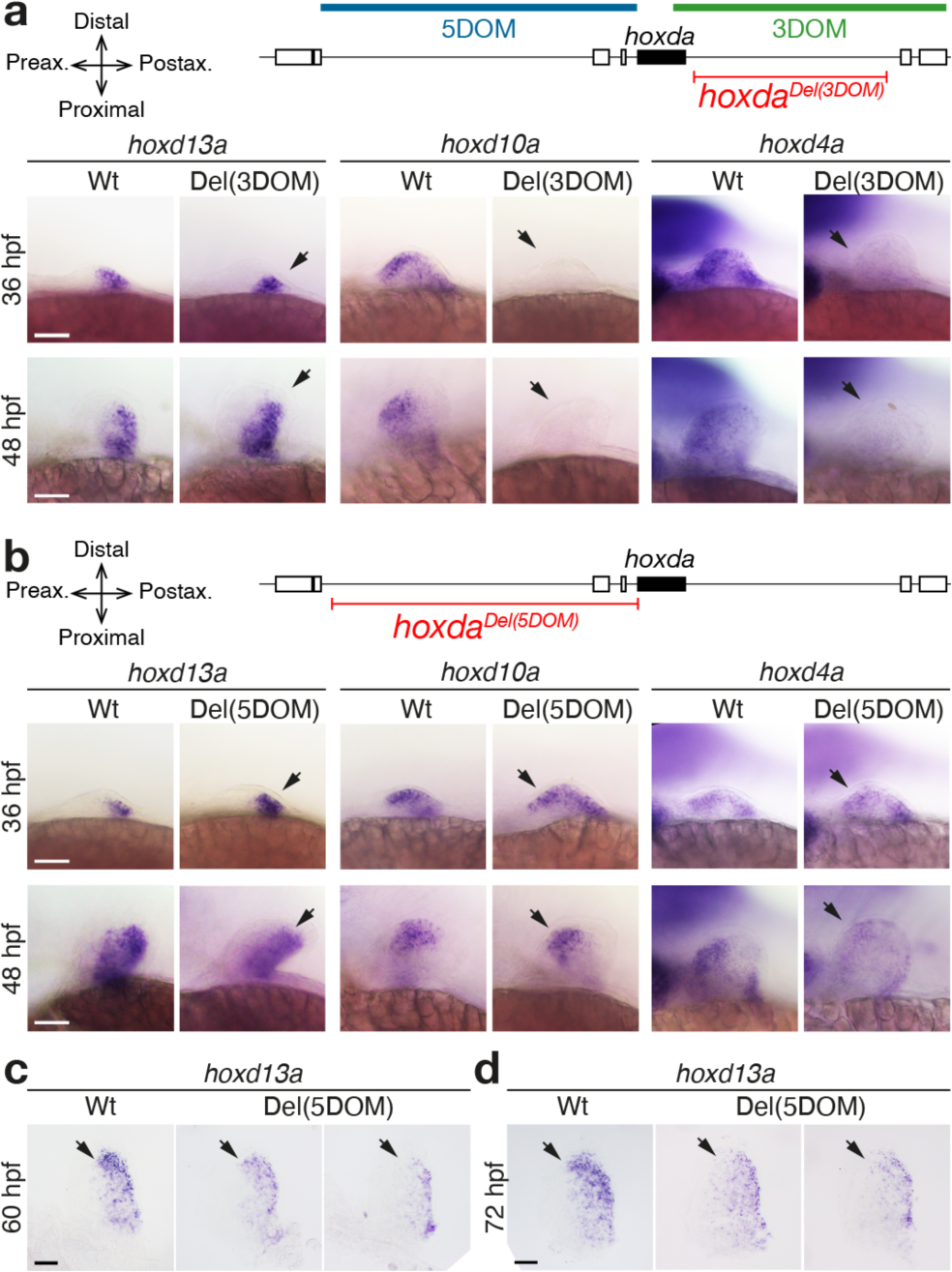
Regulation of *hoxda* genes in pectoral fins lacking the 3DOM and 5DOM regulatory landscapes. *hoxd13a*, *hoxd10a* and *hoxd4a* expression by WISH at 36 hpf, 48 hpf, 60 hpf and 72 hpf in zebrafish embryos with either the 3DOM (**a**) or the 5DOM regulatory landscapes deleted (**b-d**). Wild-type and homozygous mutant embryos derived from the same cross and are shown side by side. Scale bars = 50 µm. **a.** Expression of both *hoxd10a* and *hoxd4a* is completely lost in mutant fin buds lacking 3DOM (arrows), whereas expression of *hoxd13a* is identical to that of wild-type embryos (arrows). **b.** In fin buds lacking 5DOM, expression of all three *hoxd13a*, *hoxd10a* and *hoxd4a* are identical to matched wild-type embryos up to 48 hpf (arrows). However, at 60 hpf (**c**) and 72 hpf (**d**), a clear decrease in intensity is observed throughout, yet particularly marked in the distal aspect of the fin bud (arrows). The two examples shown here are amongst the fin buds with the greatest reduction in expression (see Fig. S5).

To determine whether *hoxd13a* transcription was controlled by enhancers present within 5DOM, we similarly analyzed Del(5DOM) zebrafish embryos by WISH. Consistent with regional control of *Hox* gene transcription, neither *hoxd4a* nor *hoxd10a* expression were affected in mutant Del(5DOM) fin buds (Fig. 1b, arrowheads). Unexpectedly, though, *hoxd13a* transcripts were unaffected, with a pattern closely matching that of control fin buds (Fig. 1b, arrowheads; Fig. S5). This was surprising as the entire regulation required for *Hoxd13* expression in tetrapods is located within this region, and previous transgenic results using components of the fish 5DOM sequences in transgenic mice showed sufficiency to drive expression^12,19^. At later stages of development, however, an effect of the 5DOM deletion on *hoxd13a* expression is suggestive, though variable. Yet *hoxd13a* expression remains globally similar to wildtype (Fig. 1c, d; Fig. S5).

These two genomic regions also control expression in other axial systems in mice^23,24^. Thus, we extended our analysis to assess shared components of regulation between these regulatory landscapes. Mutant Del(3DOM) embryos did not reveal visible differences in expression from controls in the trunk (*hoxd13a, hoxda10a, hoxd4a*), the pseudo-cloacal region (*hoxd13a*) or the branchial arches and rhombomeres (*hoxd4a*) (Fig. S6a). Del(5DOM) embryos also showed comparable expression to wildtype controls, except for the complete disappearance of *hoxd13a* transcripts from the pseudo-cloacal region (Fig. 2). We noticed a temporary reduction of *hoxd13a* expression in the tailbud (Fig. 2a), yet this deficit was no longer detectable at 36hpf. These results reveal that in zebrafish, 5DOM-located enhancers regulate *hoxd13a* genes in the cloacal area, from its onset of expression until at least 72hpf, whereas neither *hoxd10a,* nor *hoxd4a* are expressed there (Fig. S6). As previously reported for both *hoxd13a* and *hoxa13b*^25,26^, these transcripts were found in this region within of the nascent pronephric duct and hindgut. These structures eventually converge towards a single pseudo-cloacal complex that exits the body at adjacent openings without ever completely fusing. In 72hpf larvae, *hoxd13a* mRNAs also appeared in the posterior gut in both control and mutant samples, however still absent in the mutant cloacal region (Fig. 2c, black and red arrows, respectively), indicating that these two expression specificities are regulated separately.

**Figure 2.**
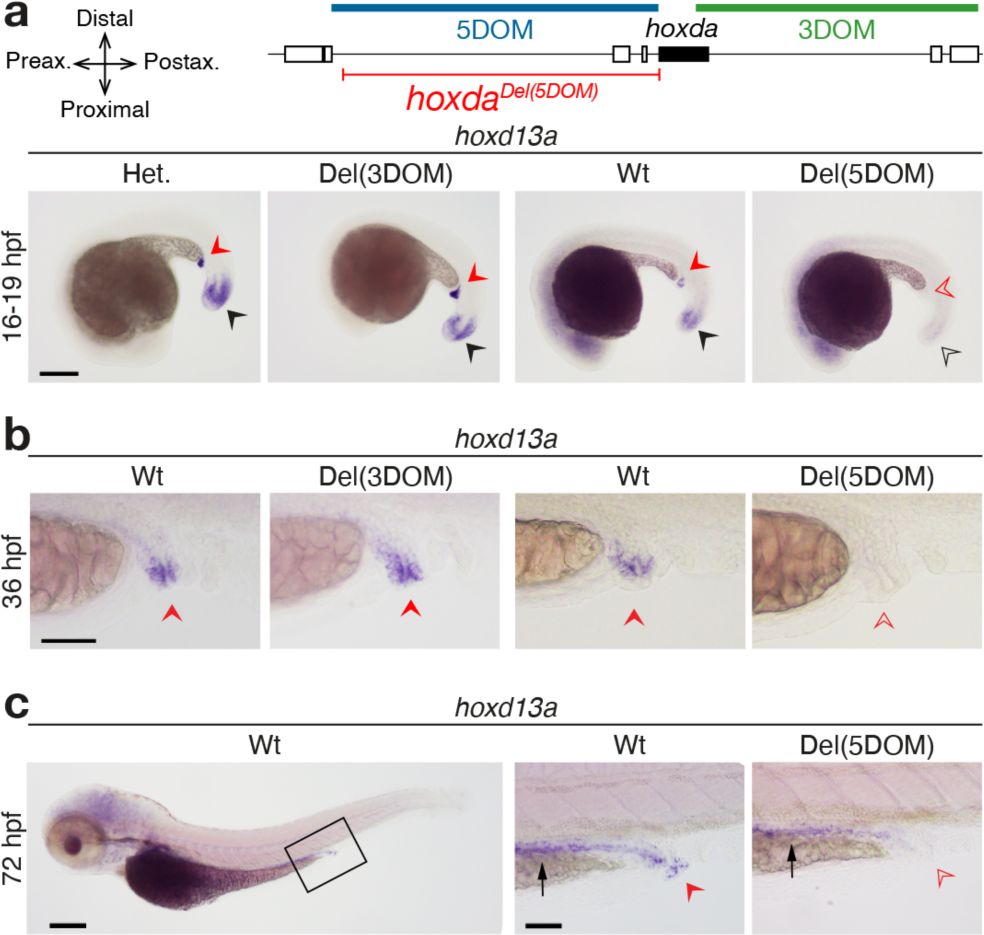
Effects of deleting 5DOM on *hoxd13a* regulation in the pseudo-cloacal region. **a-c.** Expression of *hoxd13a* is completely lost in the cloaca of 16 hpf (**a**), 36 hpf (**b**) and 72 hpf (**c**) embryos lacking 5DOM (red arrowheads), while it is identical to controls in embryos lacking 3DOM (red arrowheads), indicating that the 5DOM is required for *hoxd13a* activity in the pseudo-cloacal region. At 16 hpf, a temporary decrease of *hoxd13a* expression in the tailbuds lacking 5DOM (black arrowheads), but this effect was no longer observed at later stages. **b.** Enlargements of the cloacal region showing *hoxd13a* transcripts mostly lining the very end of the intestinal canal, converging towards the cloacal region. **c.** At 72 hpf, *hoxd13a* expression is detected in the posterior epithelial part of the gut in both control and mutant larvae (black arrow), indicating that expression in the cloacal region (red arrow) responds to a separate regulatory control. Scale bars: 200 µm for whole embryos, 50 µm for zoomed-in views.

The cloaca evolved at the base of the craniate lineage as a single orifice for the digestive, urinary and reproductive tracts, as found in birds or squamates. In mammals, a cloaca initially forms early on during embryonic development, but as the embryo grows, it divides into different openings for the urogenital and digestive systems. To evaluate whether the observed 5DOM regulation of *hoxd13a* in the zebrafish pseudo-cloacal region is a derived or an ancestral condition, we looked at the developing mouse urogenital sinus (UGS), a structure that derives from the mammalian embryonic cloacal area.

### *Hoxd* gene expression and regulation in the urogenital sinus

The UGS, positioned below the urinary bladder, is derived from a cloacal rudiment originating from hindgut and ectodermal tissue^27,28^. During mid-gestation, as the nephric and Müllerian ducts grow towards the posterior end of the embryo, they meet and fuse with the invaginating cloaca. We performed WISH on dissected urogenital systems from control murine male and female embryos at E18.5 (Fig. 3a, b). All genes tested but *Hoxd13* were detected in the anterior portions of the urogenital system including the kidneys, uterus, and deferens ducts^29,30^ (Fig. S7a, b). In contrast, *Hoxd13* expression was restricted to the UGS in both male and female embryos, along with *Hoxd12*, *Hoxd11* and, to a weaker extent, *Hoxd10* (Figs. 3 and S7), i.e., the same four genes responding to both the digit and external genitals long-range regulations exerted by 5DOM^31^, thus suggesting a transcriptional control coming from this same 5’-located domain.

**Figure 3.**
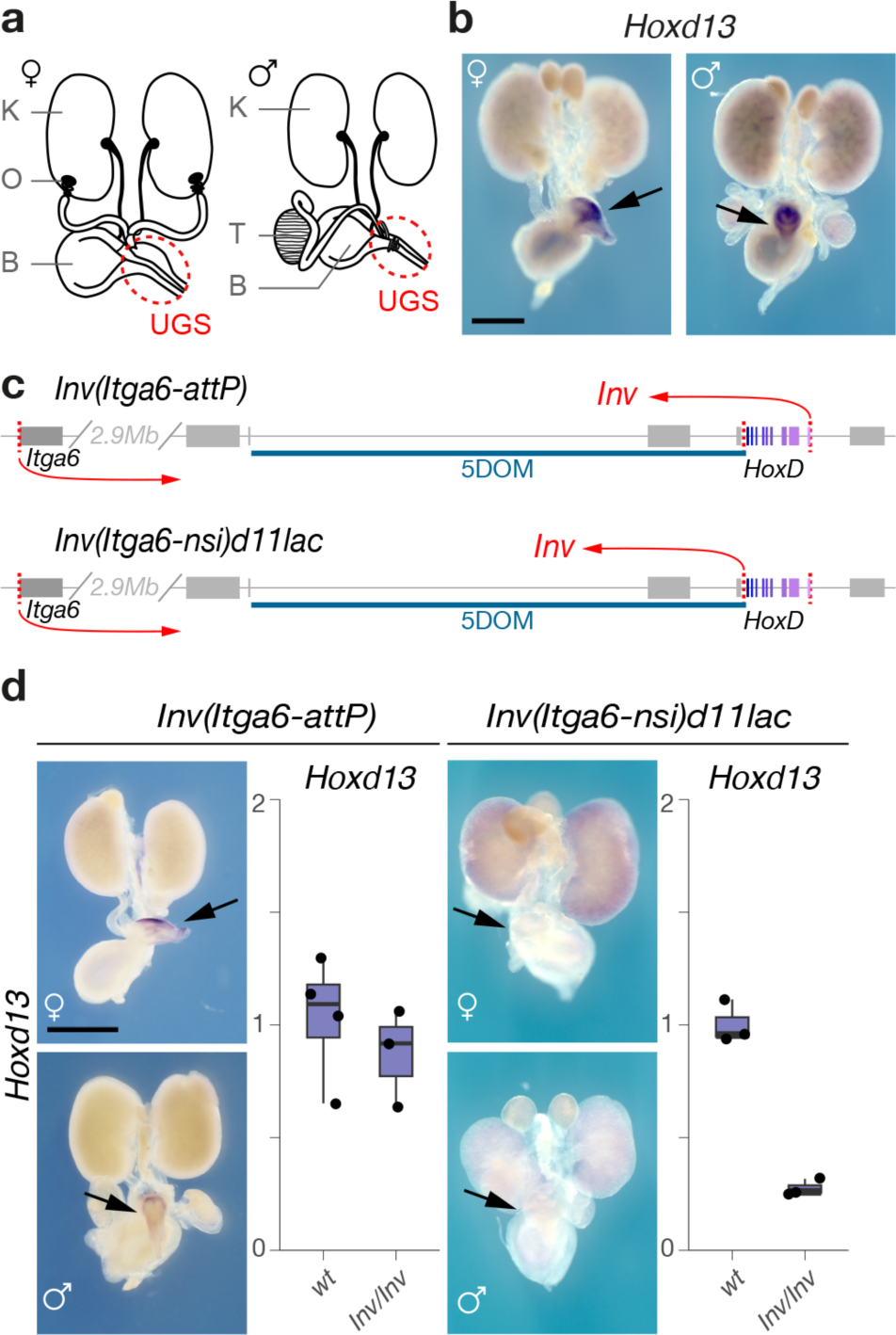
*Hoxd* gene expression in the mouse urogenital system. **a.** Schematic representations of male and female urogenital systems. K: Kidney, B: Bladder, O: Ovary, T: Testis. The urogenital sinus (UGS) is indicated with a red circle. **b.** WISH of *Hoxd13* in representative female and male urogenital systems. *Hoxd13* is selectively expressed in the UGS. **c.** Schematic representation of the two *HoxD* inversion alleles. The locations of the inversion breakpoints are depicted with red arrows. *Hox* genes shown in shades of purple. **d.** *Hoxd13* expression in urogenital systems of mice carrying the inversions (WISH, left panel; RT-qPCR, right panel). Expression of *Hoxd13* in the UGS is abolished when the target genes are disconnected from 5DOM. Scale bars: 1 mm.

We verified this by using an engineered inversion that keeps the *HoxD* cluster linked with 5DOM, but takes them far away from 3DOM (Fig. 3c, *HoxD^Inv^*(Itga6–AttP))^32^. In this allele, *Hoxd13* transcription in UGS was unaffected (Fig. 3d). We then tested a comparable inversion, yet with a breakpoint immediately 5’ the *HoxD* cluster thus disconnecting 5DOM from all *Hoxd* genes (Fig. 3c, *HoxD^Inv^*(Itga6–nsi)*^d^*^11l*ac)*^)^33^. We scored a virtually complete loss of *Hoxd13* transcription (Fig. 3d), suggesting that most, if not all, UGS-specific enhancers are located within 5DOM. We confirmed this by using a large BAC transgene covering the entire *HoxD* cluster (Fig. S7c, d)^32^ introduced into mice lacking both copies of the *HoxD* locus^34^ (Fig. S7c). In this mutant allele, *Hoxd13* transcription was not detected (Fig. S7c, arrows). Finally, we looked at beta-gal staining of a *LacZ* reporter integrated into the BAC transgene. While the reporter was strongly active in fetal kidneys as expected^24^, the UGS was not stained (Fig. S7d). In contrast, a *LacZ* reporter transgene integrated within the inversion separating 5DOM from the *HoxD* cluster (Fig. S7d, *HoxD^Inv^*(Itga6–nsi)*^d^*^11l*ac)*^) robustly stained E18.5 UGS, supporting again the presence of UGS enhancers within 5DOM (Fig. S7d).

We quantified the reduction in *Hoxd* gene expression in the *HoxD^Inv^*(Itga6–nsi)*^d^*^11l*ac)*^ allele using RNA-Seq on E18.5 UGS of males and females. In both cases, *Hoxd13*, *Hoxd12* and *Hoxd10* transcription levels dropped abruptly when compared to wildtype samples, while the transcription level of other *Hoxd* genes was not affected (Fig. S8a). Altogether, this allelic series demonstrated that the mammalian 5DOM contains the UGS enhancers, similar to the zebrafish 5DOM containing cloacal enhancers. It also showed that *Hoxd* genes responsive to this regulation (*Hoxd13*-*Hoxd10*) are the same sub-group that responds to both digit and external genitals enhancers.

### Identification of UGS enhancers

To identify UGS enhancers within the mouse 5DOM, we used three scanning deletion alleles covering 5DOM^2^ (Fig. 4a, red) and measured the change in expression by RTqPCR (Fig. S8b). In the *HoxD^Del^*(Atf2–SB1) allele, the most distal portion of 5DOM was removed with no impact on *Hoxd* gene expression levels. However, when either the central (*HoxD^Del^*(SB1–Rel5)) or the most proximal (*HoxD^Del^*(Rel5–Rel1)) portions of 5DOM were removed, transcription of *Hoxd13*, *Hoxd12* and *Hoxd10* were significantly reduced indicating that these two 5DOM intervals contain UGS enhancers (Fig. S8b).

**Figure 4.**
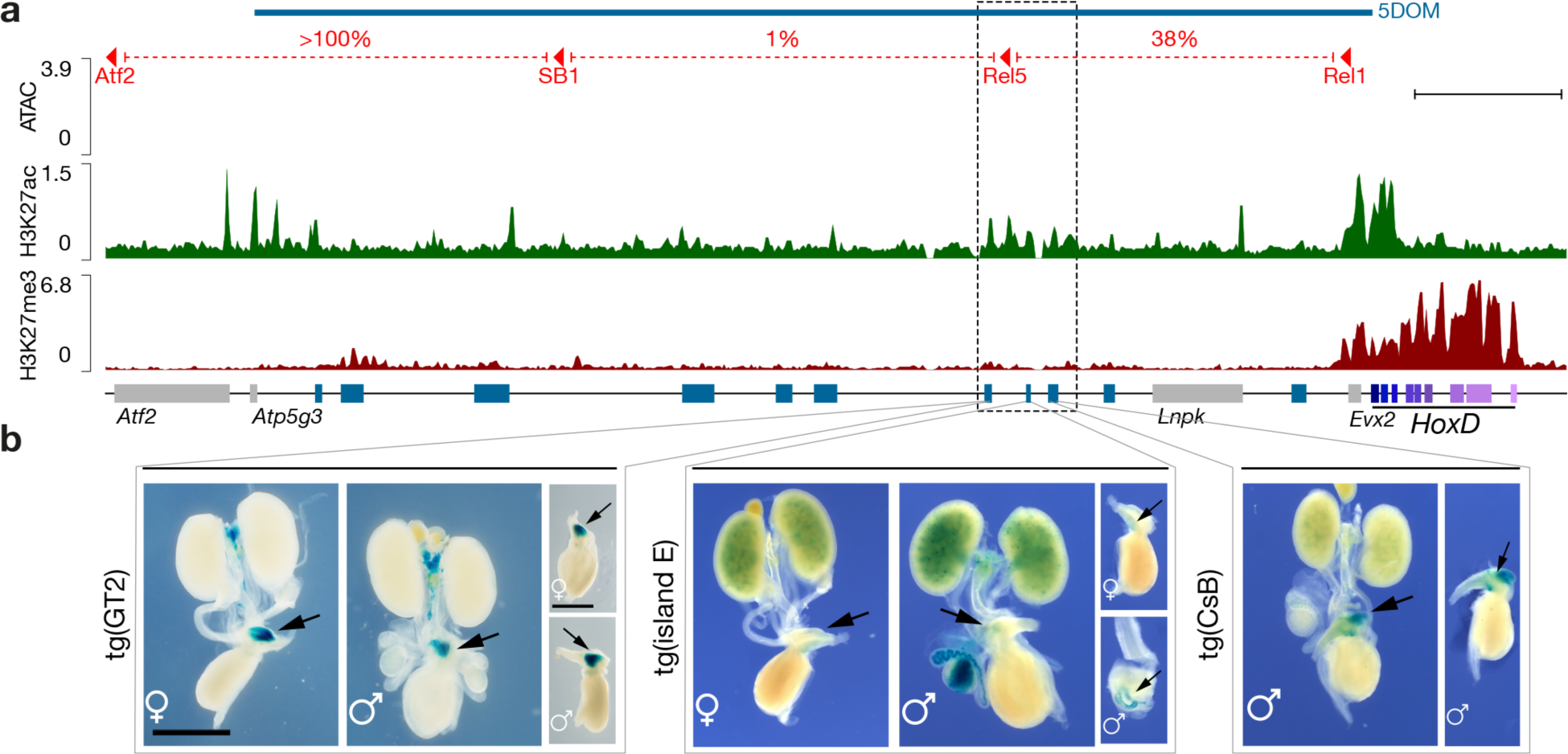
Urogenital sinus enhancers located in the 5DOM. **a.** Chromatin accessibility (ATAC-seq, blue track) and H3K27ac (green track) and H3K27me3 (red track) ChIP-seq profiles from micro-dissected male UGS at E18.5. The red lines on top delineate the three deletions within 5DOM with the percent of *Hoxd13* expression left in the UGS after each deletion (see also Fig. S8). *Hoxd* genes are in purple. Blue rectangles indicate previously described 5DOM enhancers. The dashed box highlights a H3K27ac-positive cluster of ATAC-seq peaks lacking H3K27me3 and containing three enhancer sequences; GT2, island E and CsB. Scale bar; 100 kb. **b.** Regulatory potential of the GT2, island E and CsB elements when cloned into a *lacZ* reporter cassette. GT2 induces robust *lacZ* expression in the UGS of both male and female embryos, while island E shows weaker expression. The CsB transgene induces robust expression in males (no data available for females). Scale bar: 1 mm.

We then measured chromatin accessibility by ATAC-seq and profiled H3K27ac and H3K27me3 histone marks associated with either active or inactive chromatin, respectively, by ChIP-seq on micro-dissected male UGSs (Fig. 4a). We identified a cluster of several conspicuous ATAC-seq and H3K27ac signals located approximately 200 kb upstream *Hoxd13,* in a region encompassing the Rel5 breakpoint, i.e., between the Del(SB1-Rel5) and the Del(Rel5-Rel1) deletions (Fig. 4a, dashed box). Within this 67 kb-large region, the ATAC and H3K27ac signals matched three elements previously characterized as enhancer sequences, yet with distinct tissue specificities; The GT2 and Island E sequences had been identified as a pan- and a proximal-dorsal genital tubercle specific enhancers, respectively^35,36^, whereas the CsB element was reported as a distal limb and fin enhancer element^1,12^. The ATAC peaks were positioned at regions that are relatively depleted for the H3K27ac mark (Fig. S9), which is a hallmark of active enhancer elements^21^.

For the GT2 and CsB elements, portions of the region were highly conserved across bony fish *hoxda* loci. In contrast, Island E contained a small portion of sequence only conserved amongst mammals (Fig. S9). We tested these three putative enhancers in an enhancer-reporter assay and all three sequences were able to drive robust *lacZ* expression in the UGS, closely matching the expression of posterior *Hoxd* genes in this area in both male and female specimens (Fig. 4b), indicating that in mammals, 5DOM contains a set of enhancer elements that control the transcriptional activation of *Hoxd* genes in the UGS. These three enhancers had been previously identified as specific for either distal limbs or external genitals, i.e., two structures that depend upon the 5DOM regulatory landscape as the only source of enhancers for their development. In zebrafish, while the orthologous 5DOM landscape is indeed necessary to activate posterior *hoxda* genes in the cloacal region, expression of these genes in the developing fin is not dramatically altered.

### An ancestral regulatory landscape for an ancient function

In mammals, the combined mutation of both *Hoxa13* and *Hoxd13* has a drastic effect on the development of the posterior part of the digestive and urogenital systems^30,37^, causing an absence of any detectable UGS^30^. Previous studies revealed the expression of most *hox13* genes in the developing zebrafish intestinal and cloacal regions (Fig. S10) that is suggestive of functional conservation^38^. Supporting this, 5’ *hoxa* genes were differentially regulated in the normal patterning of the goby cloacal region^39^. Therefore, we wondered whether cloacal patterning would be a core ancestral function of *Hox13* terminal genes regulation. We thus asked what if *hox13* gene function had specific roles in cloaca formation in zebrafish.

Wild-type zebrafish exhibit a pseudo-cloacal configuration in which the hindgut and pronephric duct exit the trunk through separate but adjacent openings. The outlet of the hindgut is anterior to that of the pronephric duct, and a septum resides in between (Fig. 5a, e). Homozygous single mutants of *hoxa13a*, *hoxa13b*, and *hoxd13a* are indistinguishable from the wild-type arrangement (Fig. 5b, f), as are animals triply heterozygous for these genes (Fig. S11). However, combined *hoxa13a*;*hoxa13b* double homozygous mutants exhibit connection of the hindgut with the pronephric duct before exiting the body through a single opening (Fig. 5c, g). The loss of *Hoxa13* paralogs was also found to affect pronephric duct and hindgut length at the level of the median fin fold (Fig. S11). A more severe phenotype is observed in *hoxa13a*;*hoxa13b*;*hoxd13a* triple mutants, in which the septum is dysmorphic and the hindgut and pronephric duct are fused, resulting in a large shared lumen and outlet (Fig. 5d, h). These results reveal a conserved requirement of *Hox13* function for the normal patterning of the termini of digestive and urogenital systems across vertebrates.

**Figure 5.**
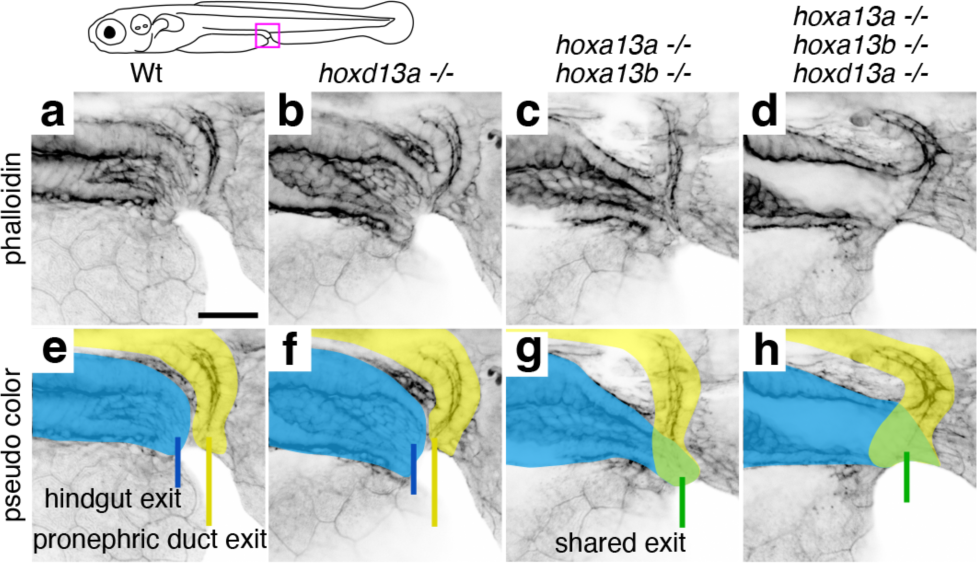
Loss of *hox13* paralogs in the zebrafish results in defects of the cloacal region. Confocal microscopy of phalloidin-labeled cloacal regions of wild-type and *hox13* mutant zebrafish at 6 days post-fertilization shown in a single channel (**a-d**) and with pseudo coloring (**e-h**). Pseudo coloring indicates hindgut (blue), pronephric duct (yellow), or fused ducts (green). **e.** Wild-type fish have adjacent but distinct openings for the hindgut (blue line) and pronephric duct (yellow line) (n= 2), as do *hoxd13a* mutants (n=2) (**f**). **g.** *hoxa13a*;*hoxa13b* double mutants exhibit fusion of the hindgut and pronephric duct and a single opening (green line) (n=2). **h.** *hoxa13a*;*hoxa13b*;*hoxd13a* triple mutants show connection of the hindgut and pronephric duct to form a large shared lumen (green) with a single opening (green line) (n=4). Scale bar: 30 μm.

## DISCUSSION

### *Hox* regulatory landscapes and the fin to limb transition

The expression and function of *Hoxd* and *Hoxa* genes have been central to hypotheses attempting to explain the evolutionary change from fins to limbs (e.g.^5,11,13,14^). By making comparisons of their complex transcription patterns across actinopterygian, chondrichthyan, and sarcopterygian fishes, various efforts have sought relate the two types of paired appendages. These analyses have led to the conclusion that, despite being composed of different types of skeletons, the development of actinopterygian fin rays and digits have a common regulatory architecture^11,40^. Here, by deleting the two TADs flanking the fish *hoxda* cluster, we show that the essential digit regulatory landscape characterized in tetrapods indeed has a structural counterpart in teleosts (Fig. S1)^18,41^. However, we report that, unlike in limbs, only a small part of the regulation controlling *hoxda* gene expression in distal fin buds is located within this landscape. Indeed, while some enhancer(s) controlling *hoxd13a* transcription in developing zebrafish fins are located within 5DOM^12^, most of the regulatory control likely resides within the gene cluster itself, probably at the vicinity of the *hoxd13a*, *hoxd12a* and *hoxd11a* genes, i.e., the three genes sharing the same expression in post-axial cells^16^.

This observation is consistent with the absence, in the zebrafish 5DOM, of sequences related to several strong mouse digit enhancers^12^, but also confirms results obtained when assaying fish 5DOM conserved sequences as transgenes, either in zebrafish or in mice^12,19^. It also explains why the zebrafish *lnpa* gene, which is embedded into 5DOM, is not expressed in the emerging pectoral fin buds^16^, whereas the mouse counterpart has a strong distal expression due to enhancer hi-jacking^1^. The presence of such a partial distal landscape in teleosts may illustrate an intermediate step in the full co-option of this regulation, as achieved in tetrapods (see below). Alternatively, it may reflect a secondary loss of several distal enhancers associated with teleost whole genome duplication. These questions may be solved with a comparable deletion and epigenetic characterization of 5DOM in more basal fish species such as gar, sturgeon, or even sharks. In contrast, the deletion of the opposite 3DOM landscape, which is responsible for all proximal *Hoxd* expression in tetrapods limb buds^42^, abolished *hoxda* gene expression in fin buds, demonstrating the genuine ancestral character of this regulation, which must have been implemented as soon as paired appendages evolved.

### An ancestral cloacal regulation

Zebrafish *hoxa13a* and *hoxd13a,* as well as other *hox13* paralogs^43^, are strongly expressed in and around the developing cloacal region^25,26^. This is an area where the extremities of both the gastro-intestinal tract and the reproductive and urinary systems come together, even though their openings remain separated, unlike in some chondrichthyan fishes or other vertebrates where the tubes coalesce into a single opening (e.g., in sharks or birds). Here we report that this pseudo-cloacal structure is disrupted in zebrafish carrying *hox13* mutant alleles, with an abnormal fusion between the intestinal and the pronephric opening thus giving rise to a single yet abnormal, cloacal opening. Likewise, the developing murine UGS expresses *Hoxd13*^29^ and double mouse *Hoxd13-Hoxa13* mutant animals had severely malformed posterior regions^30,37^, with no distinguishable UGS^30^, illustrating that the evolutionary conservation of this regulatory landscape is accompanied by shared functional effects.

We also document that, as for zebrafish, the control of murine posterior *Hoxd* genes in the cloaca is achieved by enhancers located within the 5’ located regulatory landscape, i.e., in the same genomic region that regulates expression in both digits and external genitals. In mouse, several 5DOM enhancers are somewhat versatile such as the GT2 sequence, which is both UGS and genitalia-specific^44^, whereas CsB is UGS and digit-specific^1^. Other enhancer sequences, however, seem to have kept a unique specificity such as ‘island 2’, the strongest *Hox* digit enhancer identified thus far^45^, which is located in a different area of 5DOM^2^ and absent in zebrafish^12^. These observations illustrate a ‘functional adaptation’ of enhancers, which could be facilitated by a spatial proximity within the same large chromatin domain thus triggering the sharing of upstream factors (see^46^). Finally, all the regulatory specificities encoded in this 5DOM landscape control the same subset of posterior *Hoxd* genes in tetrapods (from *Hoxd13* to *Hoxd10*), suggesting that while groups of enhancers can be reutilized for new tissue types, there is a constraint on which genes they can target.

### Successive co-options of a regulatory landscape

In vertebrates, *Hox13* genes are located within a topologically associated domain (TAD) distinct from that including more anterior *Hox* genes and their regulations^15,47^. This condition prevents *Hox13* to be activated too early and hence too anteriorly in the body axis, a situation detrimental for the embryo due to the potent posteriorizing function of these proteins^48^. As a result, *Hoxd13* was likely the main target gene that triggered and stabilized the various evolutionary co-options of 5DOM regulations due to its location within the 5DOM TAD and through its function to organize posterior or distal body parts together with its *Hoxa13* paralog^7,30,37^. Our results indicate that the initial functional specificity of this regulatory landscape was to organize a cloacal region, which is the posterior part of the intestinal and urogenital systems. This conclusion is supported by the documented expression of *Hox13* paralogs in the cloacal regions of paddlefish^38,49^, catshark^49,50^ and lampreys^51^, suggesting this patten was a characteristic of the common ancestor of craniate vertebrates.

While this ancestral function has been maintained throughout vertebrates, it is more difficult to infer the temporal sequence of co-options of this regulatory landscape along with the evolution of digits and external genitals (Fig. 6). However, as genitalia are late amniote specializations, it is conceivable that elaboration of digital character arose initially, suggesting a first regulatory co-option of this landscape (or part thereof) from a cloacal to a digital specificity. In support of this proposal, digits appeared in aquatic sarcopterygian fishes. These fishes do not have apparent reproductive structures to facilitate internal fertilization, and many tetrapod species also do not have external genitals. A second co-option of this now multi-functional regulatory landscape then might have occurred along with the emergence of external genitals. This latter step would have been facilitated both by the developmental proximity between external genitals and the embryonic cloacal region where posterior *Hox* genes are initially expressed^52^, and by the tight developmental relationships between amniotes limbs and genitals^29,52^. Altogether, our results indicate that the repeated redeployment of an ancient regulatory landscape, first arising along with the formation of the cloaca, serve as a foundation for the evolution and elaboration of innovations in vertebrates.

**Figure 6.**
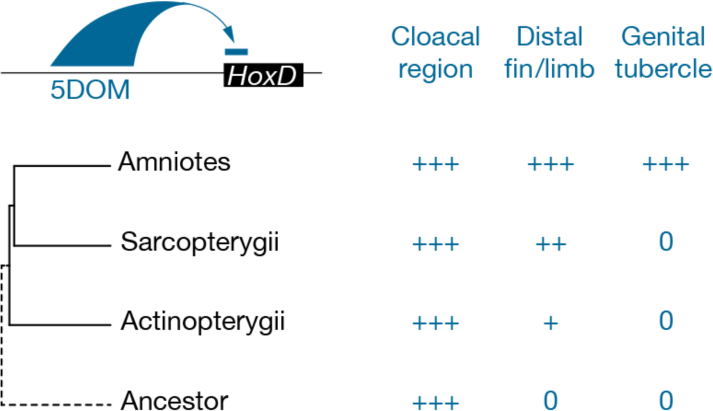
Evolutionary co-option of the *HoxD* 5DOM regulatory landscape. Schematic representation of posterior *Hoxd* gene regulation by the 5DOM regulatory landscape (top left) and the (at least) three developmental contexts where this landscape is functional (top right). On the left are shown the phylogenetic relationships between taxa where distal fins, distal limbs and external genitals emerged, while on the right, the corresponding 5DOM regulatory contributions to these structures are indicated. “0” denotes the absence of any given structure. In this view, the 5DOM cloacal regulation is an ancestral feature. In actinopterygian fishes, 5DOM lightly contributes to *hoxda* gene regulation in postaxial and distal territories of paired fin buds. The regulatory importance of the 5DOM in distal fin territories increases in sarcopterygian fishes. In amniotes, the 5DOM contribution expands to take over the entire regulation of posterior *Hoxd* genes in digits, as suggested by many enhancers with mixed specificities. Similarly, a distinct yet overlapping set of 5DOM-located enhancers entirely control *Hoxd* gene expression in the genital tubercle.

## LEGENDS TO SUPPLEMENTARY FIGURES

**Figure S1.**
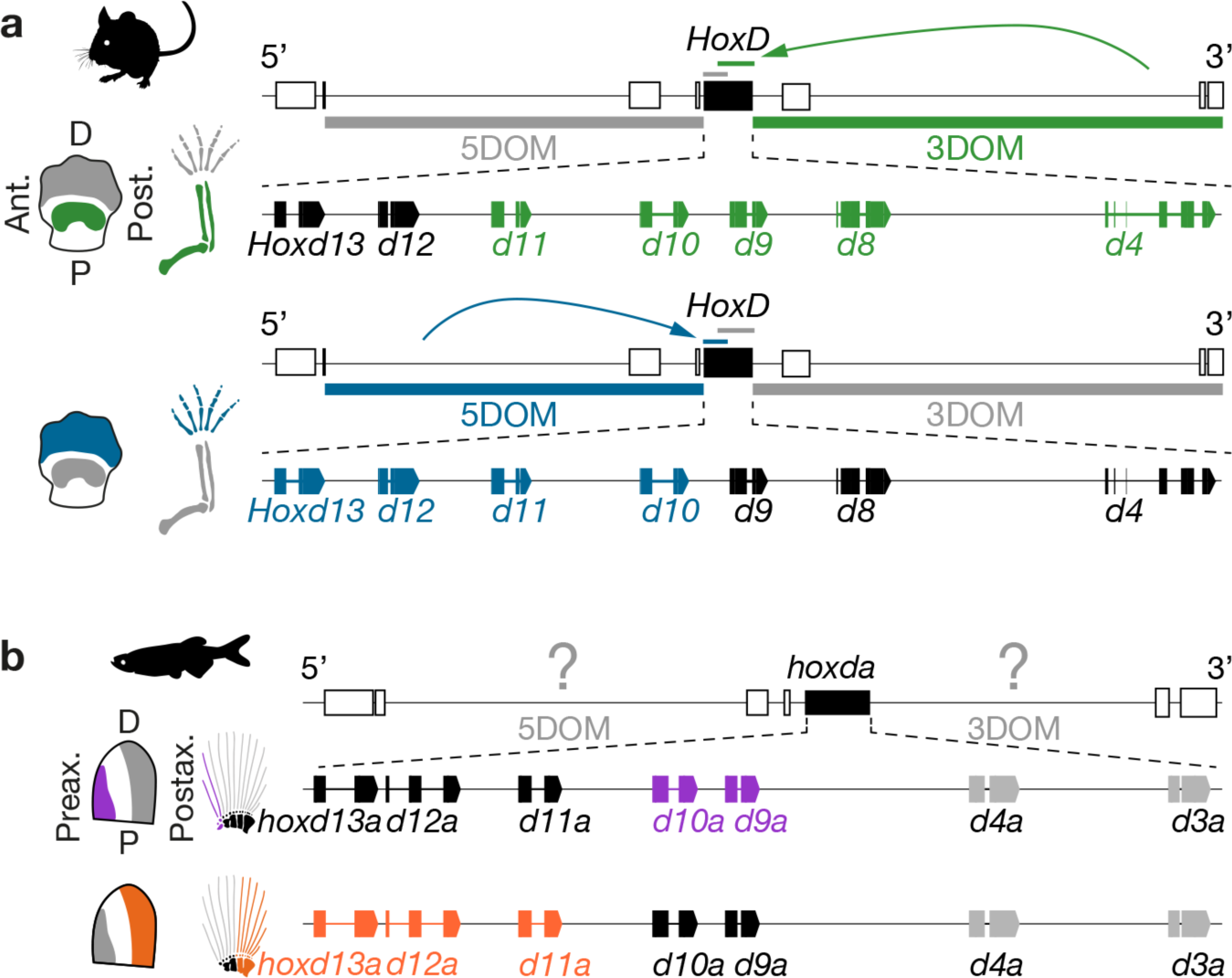
Comparison of *HoxD* regulatory landscapes in mammals and fishes. **a.** *Hoxd* gene expression and regulation in mouse limb buds at E12.5. The *HoxD* cluster is flanked by two gene deserts, named according to their relative position (3’ or 5’) with respect to *Hoxd* gene orientation. The 3DOM regulatory landscape activates *Hoxd4* to *Hoxd11* in the proximal limb territory (green). The 5DOM activates *Hoxd10* to *Hoxd13* in the distal limb territory (blue). Schemes are based on ref.^15^. **b.** Gene expression in fin buds at 40-60 hpf in the cognate zebrafish *hoxda* cluster. The fish cluster is also flanked by two gene deserts but their regulatory potentials are unknown (question marks). Fish *hoxd9a* to *hoxd11a* are expressed in the preaxial fin territory (purple) whereas *hoxd11a* to *hoxd13a* are expressed in a postaxial domain (orange). Schemes and WISH are inspired from^14,53,54^.

**Figure S2.**
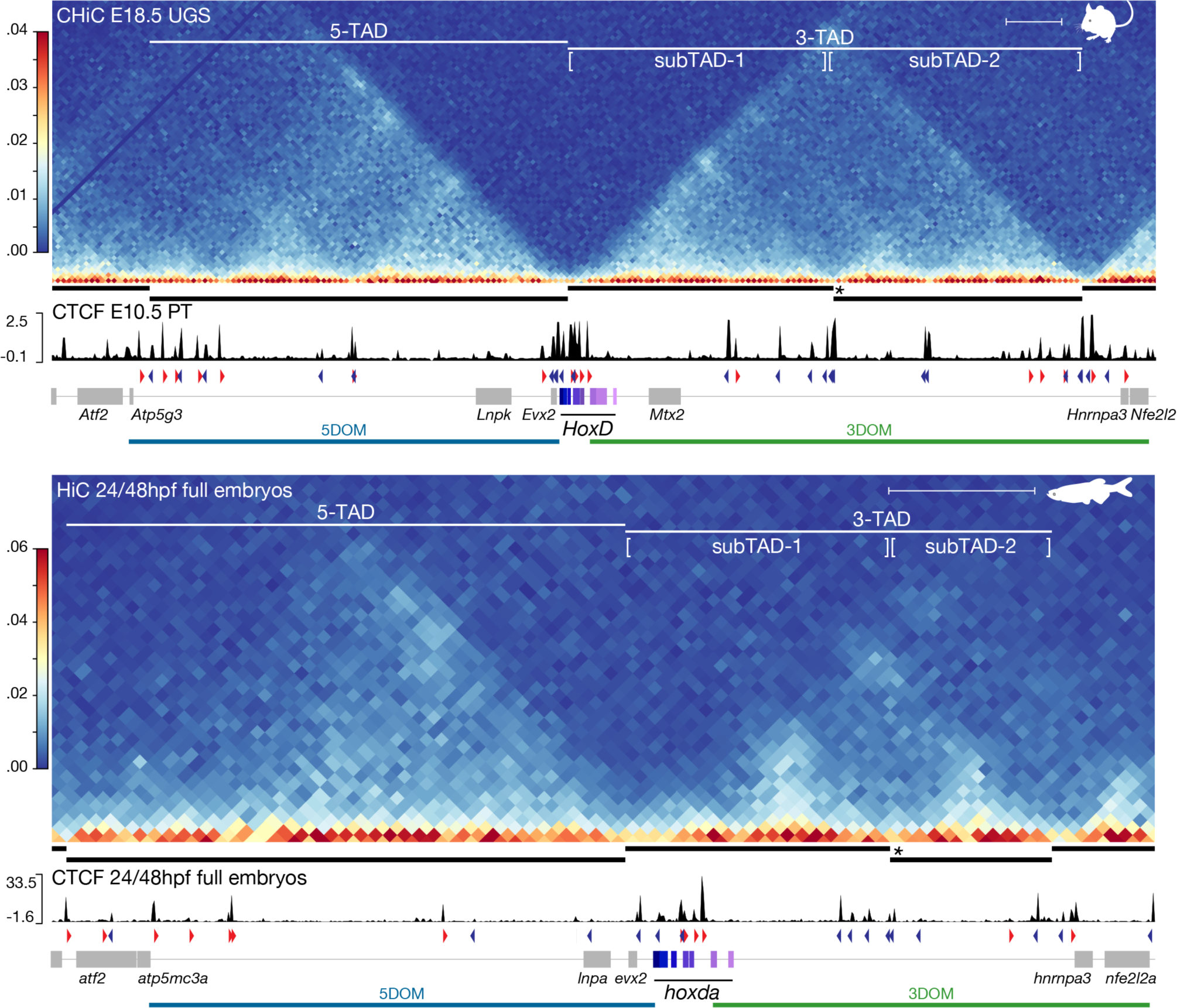
3D chromatin conformation at the mouse and fish *HoxD* loci. Contact frequency heatmaps at the mouse *HoxD* (E18.5 male UGS, one representative replicate out of two) and fish *hoxda* (24 hpf and 48 hpf total embryos^41,55^) loci (top and bottom, respectively). The similarities in the constitutive structural organization of the mouse and the fish loci are underlined either by the position and relative extents of TADs (thick black lines), the presence of a sub-TAD boundary within 3DOM (asterisk), as well as by the positions and orientation of CTCF binding sites (red and blue arrowheads). *Hox* genes are in purple-scale rectangles and other genes are grey rectangles. Bin size is 10 kb. The scales on the *x* axes were adjusted to comparable sizes for ease of comparison, yet the fish locus is more compact. Scale bars in both cases; 100 kb.

**Figure S3.**
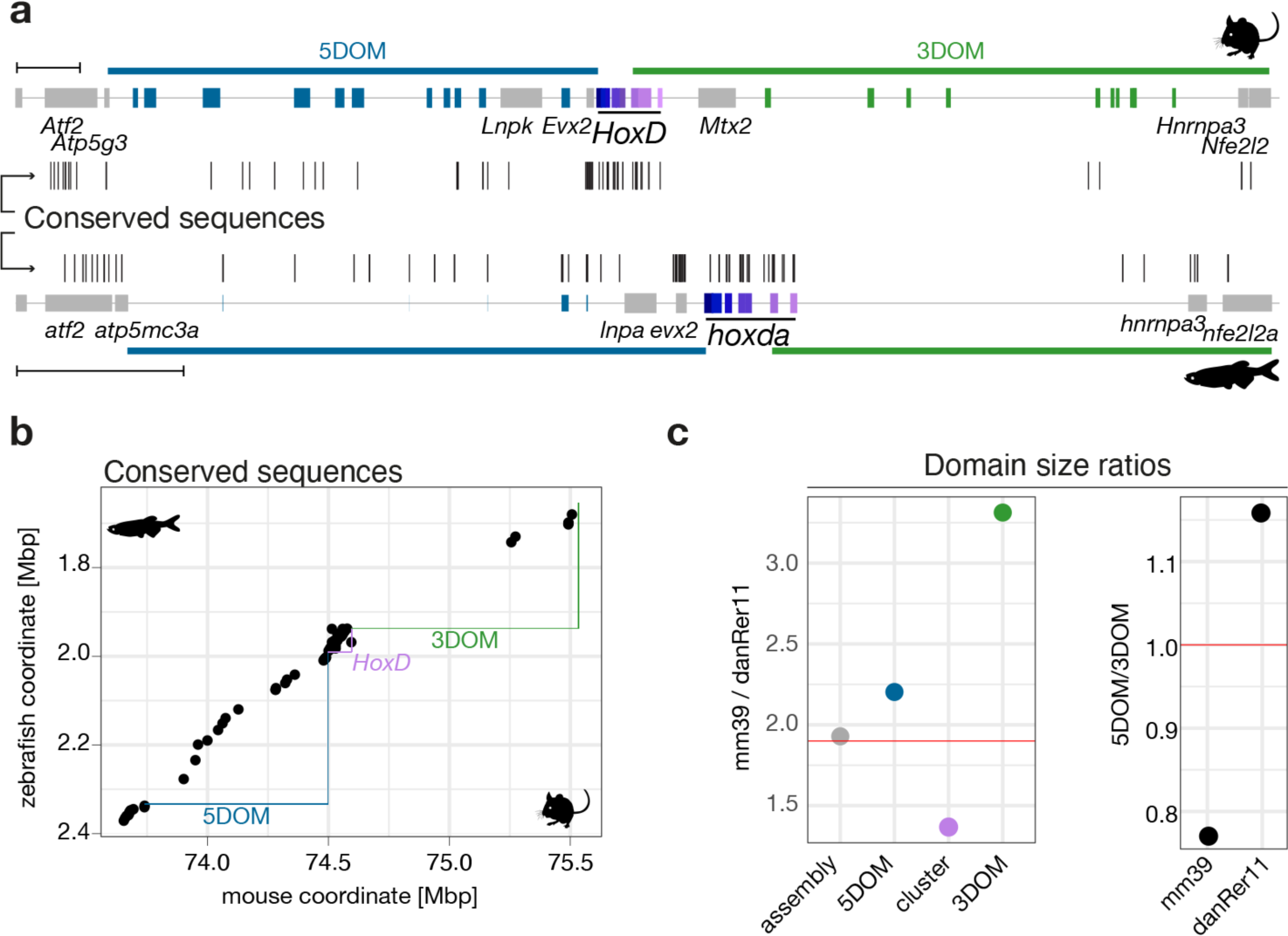
The *HoxD* locus is part of a large syntenic interval. **a.** The mouse *HoxD* locus (mm39) is on top and the zebrafish *hoxda* locus (danRer11) is shown below. *Hox* genes are **in** purple-scale rectangles and annotated mouse enhancers are shown as either blue (5DOM) or green (3DOM) rectangles. Conserved sequences between the two gene deserts are shown as vertical black bars. Those conserved sequences overlapping with known murine enhancers were used to annotate the corresponding elements in zebrafish (blue rectangles). **b.** Synteny plot representing sequences conserved between the mouse and the zebrafish *HoxD* loci. On the *x* axis is the mouse locus (mm10, chr2: 73605690-75662521) and on the *y* axis is the zebrafish locus (danRer11, chr9: 1639965-2393397, inverted *y* axis). Despite a mouse locus that is in average 2.6 times larger than its zebrafish counterpart, the order of most conserved sequences is maintained, showing the absence of substantial genomic rearrangement at these gene deserts. **c.** Size comparisons between different regions of the zebrafish *hoxda* and the mouse *HoxD* loci. The left panel shows that the *Hox* clusters have maintained a similar size over time, while gene deserts have expanded in mouse and/or contracted in zebrafish. The right panel shows that the ratio between the sizes of 5DOM *versus* 3DOM is inverted in the two species.

**Figure S4.**
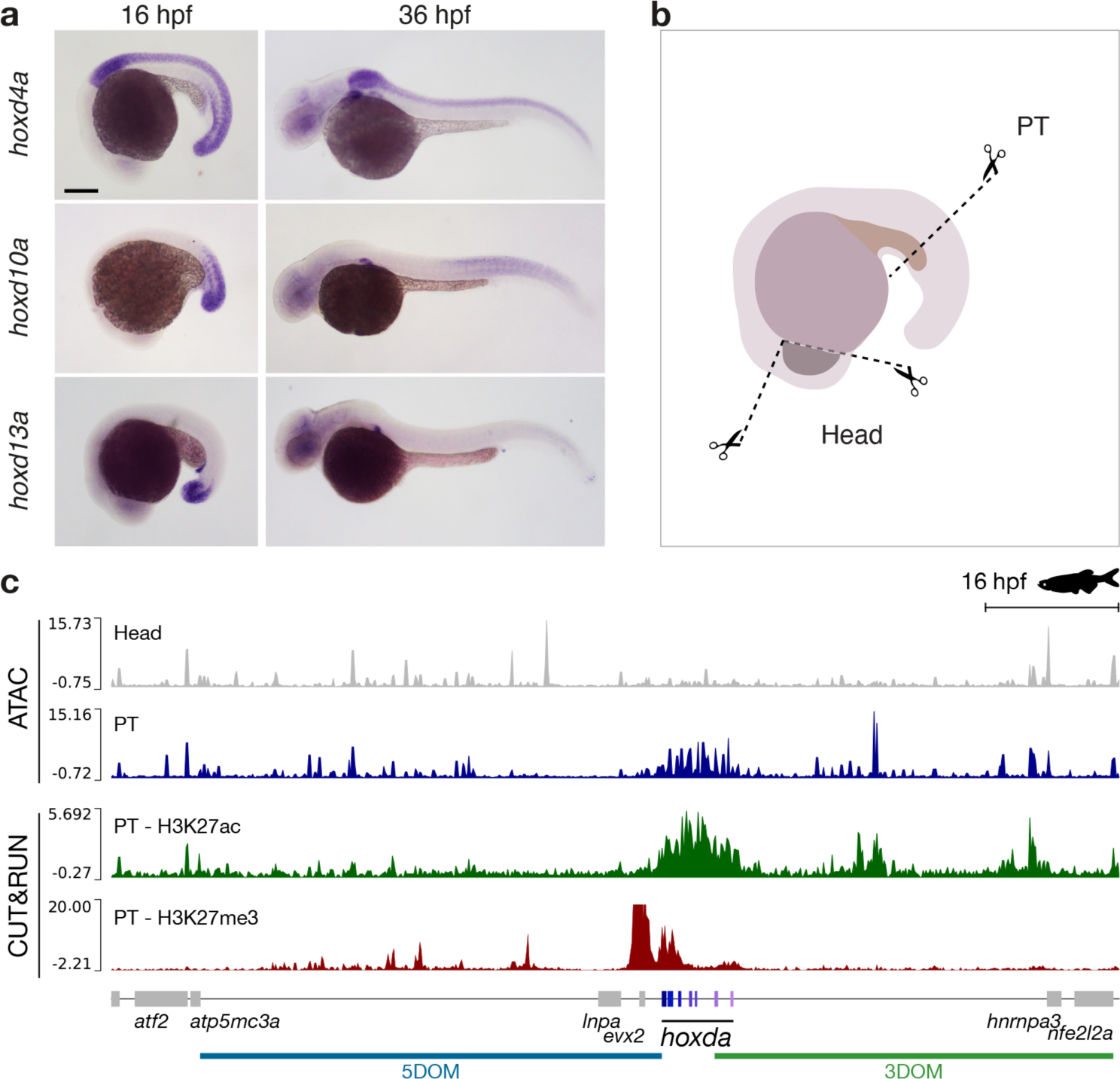
Chromatin profiles in zebrafish embryos. **a.** Expression of *hoxd13a*, *hoxd10a* and *hoxd4a* in control zebrafish embryos by WISH. Stages are indicated on top of the panels. Scale bars; 200 µm. **b.** Dissection plan used for panel (**c**). PT, posterior trunk. **c.** ATAC-seq profile and both H3K27ac and H3K27me3 CUT&RUN profiles over the zebrafish *hoxda* locus in 16 hpf dissected heads (grey, one representative condition out of three) and 16 hpf posterior trunk cells (PT, blue, one representative replicate out of three). Both the *hoxda* cluster and 3DOM show specific open sites in posterior trunk, where *hoxda* genes are expressed, when compared to forebrain. The CUT&RUN profiles in posterior trunk cells show enrichment for H3K27ac (green coverage) on the central and anterior parts of the *hoxda* cluster, while H3K27me3 (red coverage) is enriched on the posterior part and on *evx2*. Scale bar; 100 kb.

**Figure S5.**
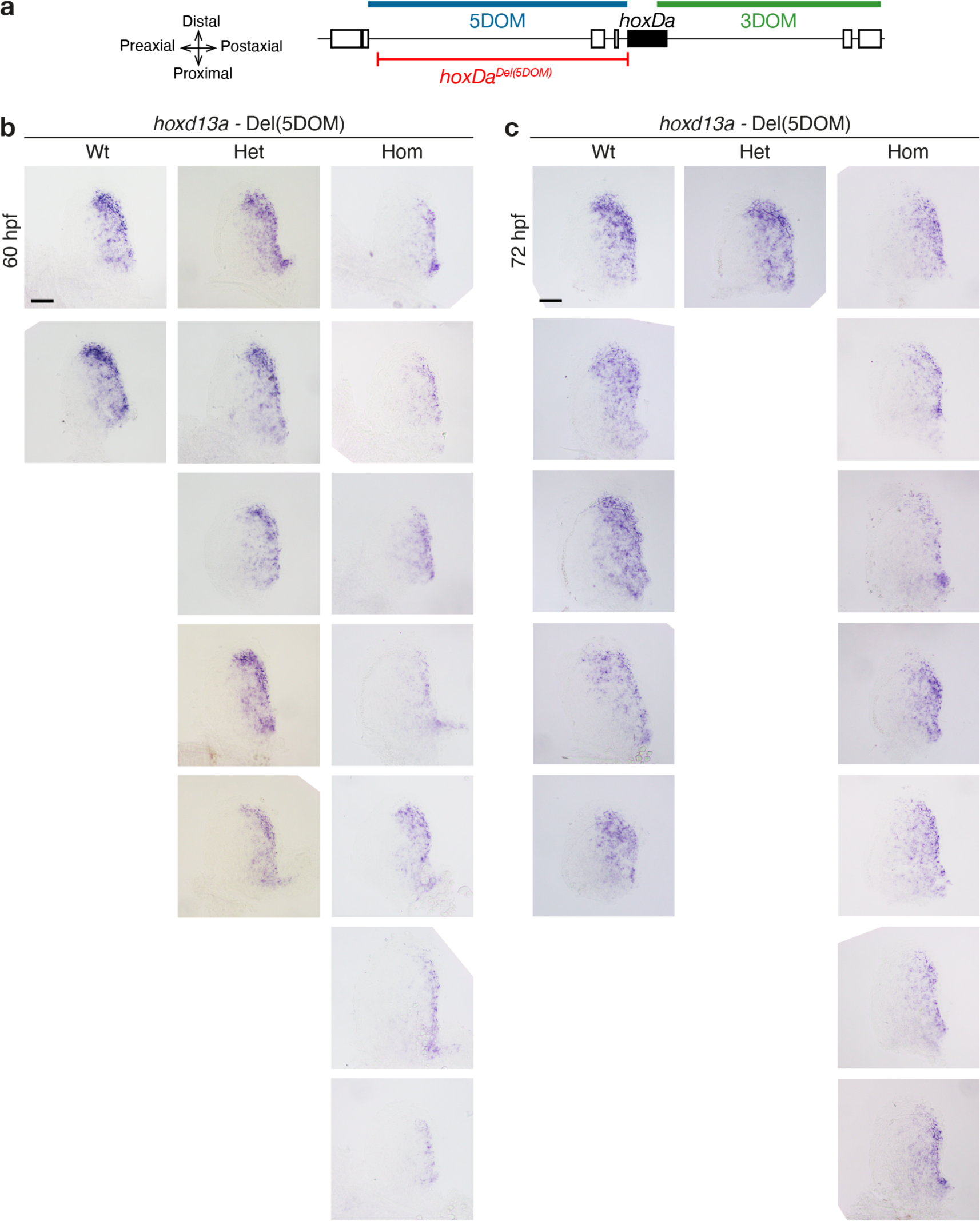
*Hoxd13a* expression in control, heterozygous and homozygous mutant fin buds at 60 and 72 hpf. **a.** Schematic of the deletion and spatial orientation of the fin buds**. b.** Various samples are shown to illustrate the variability observed. While a clear tendency is observed in the loss of the distal most expression in homozygous mutants, expression is still observed in some samples as well as in post-axial cells, unlike the situation in developing limb buds where expression is entirely absent in the comparable deletion. Scale bar: 50 µm.

**Figure S6.**
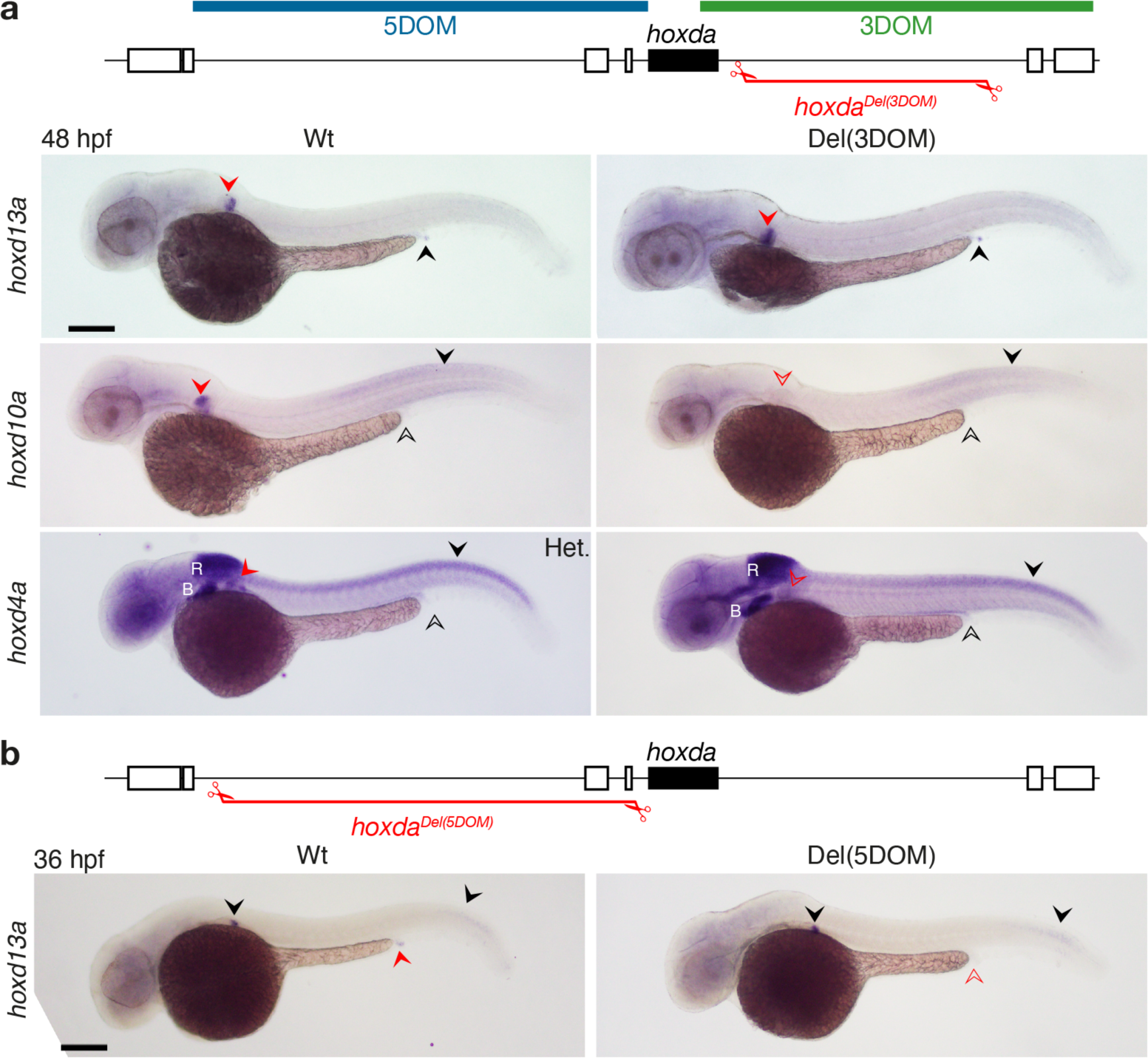
WISH of *hoxd13a*, *hoxd10a* or *hoxd4a* in zebrafish embryos lacking either 3DOM (a), or 5DOM (b). The genotypes (in red, top) and genes analyzed (left) are shown as well as the stages (up left). **a.** Deletion of 3DOM. Black arrowheads (empty for no expression and full for expression) indicate differential gene expression in the cloacal region, whereas red arrowheads (empty for no expression and full for expression) point to the pectoral fin buds. Control and homozygote mutant embryos are shown side by side for each condition, except for *hoxd4a* where a heterozygous (Het) mutant is shown. Wild-type and homozygous embryos originate from the same clutch of eggs and were processed together. In Del(3DOM) mutant embryos (**a**), *Hoxd10a* and *hoxd4a* transcription is lost in fin buds, whereas *hoxd13a* transcripts in the cloaca are not affected. B, branchial arches; R, rhombomeres; Scale bars; 200 µm. **b**. In contrast, *hoxd13a* mRNAs are lost from the cloacal region in Del(5DOM) mutant animals at 36 hpf (red arrowheads), while still clearly detected in the fin buds, indicating that the 5DOM is necessary for *hoxd13a* transcription in the pseudo-cloacal region.

**Figure S7.**
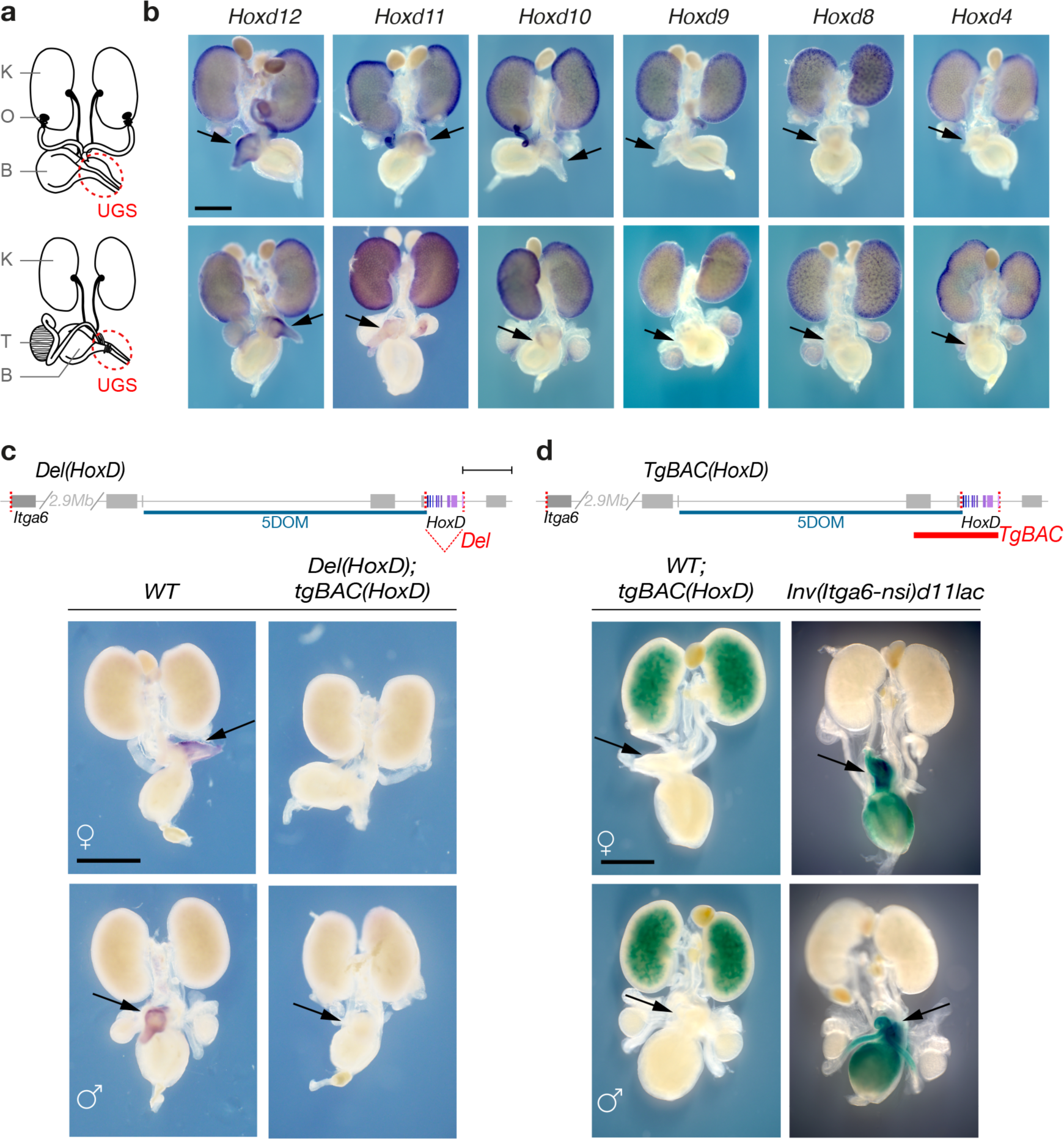
*Hoxd* gene expression in the mouse urogenital system. **a.** Schematic representations of male and female urogenital systems. K: Kidney, B: Bladder, O: Ovary, T: Testis. The urogenital sinus (UGS) is indicated with a red circle. **b.** WISH of *Hoxd* genes in representative female and male urogenital systems. All *Hoxd* genes are expressed in anterior portions of the UGS including kidneys, the uterus, deferens ducts and the bladder, except *Hoxd13* transcripts, which are restricted to the UGS (see Fig. 3b). **c-d.** Schematic representation of two *HoxD* genomic configurations, The first one is a deletion of the entire *HoxD* cluster (**c**), whereas the second one is a random integration of a transgene carrying the same *HoxD* transgene plus some flanking sequences in 5’ (**d**, thick red bar). *Hox* genes as in shades of purple and the deletion breakpoints are shown as vertical dashed red lines. Scale bar; 100 kb. **c.** WISH of *Hoxd13* in UGS from a transgenic *HoxD* cluster (TgBAC), while lacking both endogenous copies of the *HoxD* cluster. Expression is not detected from the transgenic cluster. **d.** Likewise, ß-gal staining of UGS transgenic for the *HoxD* cluster containing a *LacZ* reporter displays no activity in the UGS. By contrast, *LacZ* staining of mutant *Inv(nsi-Itga6)d11lac* embryos, which also includes a *lacZ* reporter confirms that 5DOM is necessary and sufficient to drive expression in the UGS. Scale bar: 1 mm.

**Figure S8.**
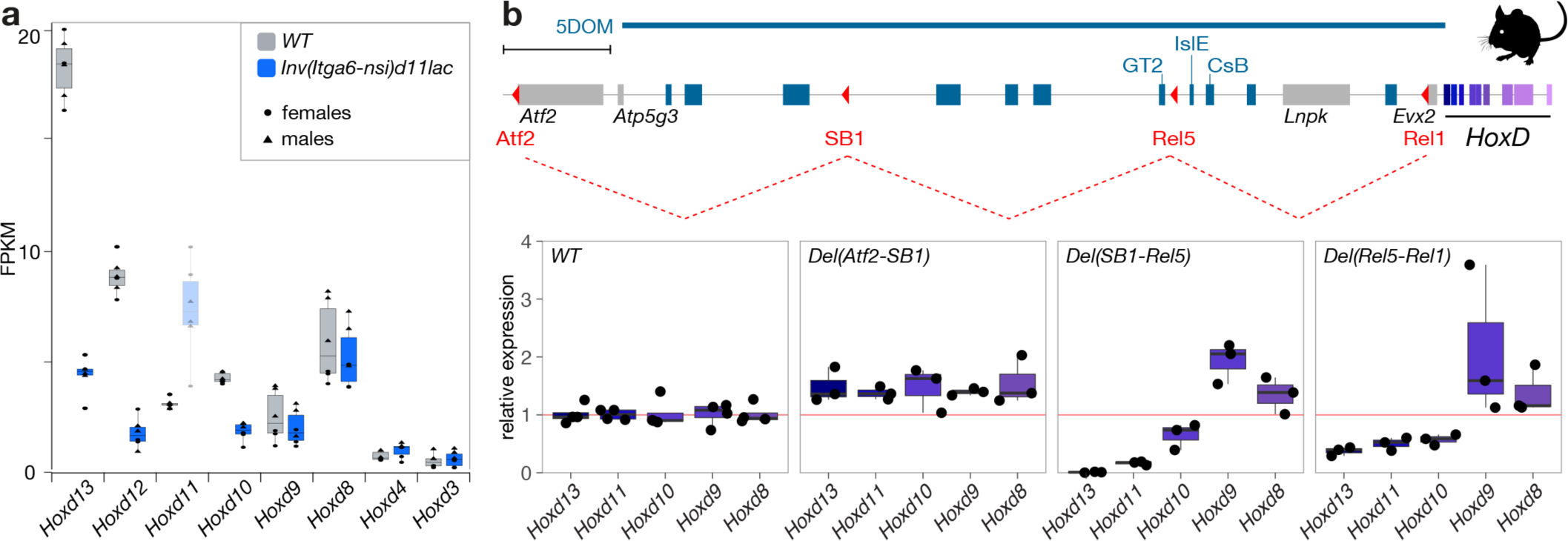
Regulatory potential of sub-regions within 5DOM. **a.** RNA-seq FPKM values for various mouse *Hoxd* genes in E18.5 UGS obtained from either wild-type or *Inv(nsi-Itga6)d11lac* mutant embryos (see schematic in Fig. 3c). Data are shown separately for females (n=3, dots) and males (n=3, triangles). Drastic decreases are observed for *Hoxd10, Hoxd12* and *Hoxd13* when 5DOM is disconnected from the *HoxD* cluster. *Hoxd11* could not be assessed due to the presence of a transgenic copy of this gene in the *LacZ* reporter cassette. **b.** On top is a scheme of the 5DOM regulatory landscape on mm39 with *Hox* genes in purple. Blue rectangles indicate previously described 5DOM enhancers. The red arrowheads delimit the serial deletion breakpoints. The three consecutive deletions are depicted by red dashed lines. Below are RT-qPCR quantifications of expression levels relative to wild-type (n=4) in three mutant lines carrying serial deletions of 5DOM (n=3). The horizontal red line represents the value of 1 for reference. Severe reductions are observed for both the *Del(Rel5-SB1)* and *Del(Rel1-Rel5)* conditions, unlike in the *Del(SB1-Atf2)* deletion. Scale bar; 100 kb.

**Figure S9.**
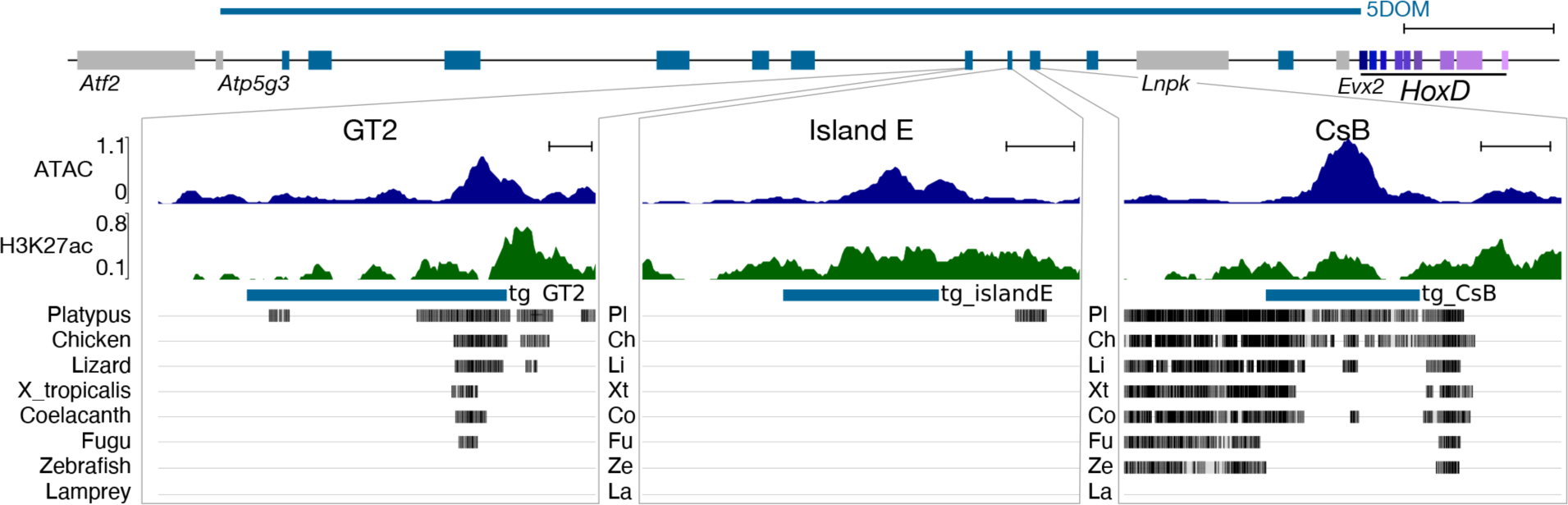
Sequence conservation in vertebrates of the GT2, islE and CsB UGS enhancers. All three sequences are comprised in the box highlighted in Fig. 4a. The ATAC-seq and H3K27ac ChIP-seq profiles are shown with, below, their sequence conservation from fishes and mammals. The thick blue lines below the H3k27ac profiles indicate the extent of the transgenes assayed in Fig. 4. Scale bars; 1 kb.

**Figure S10.**
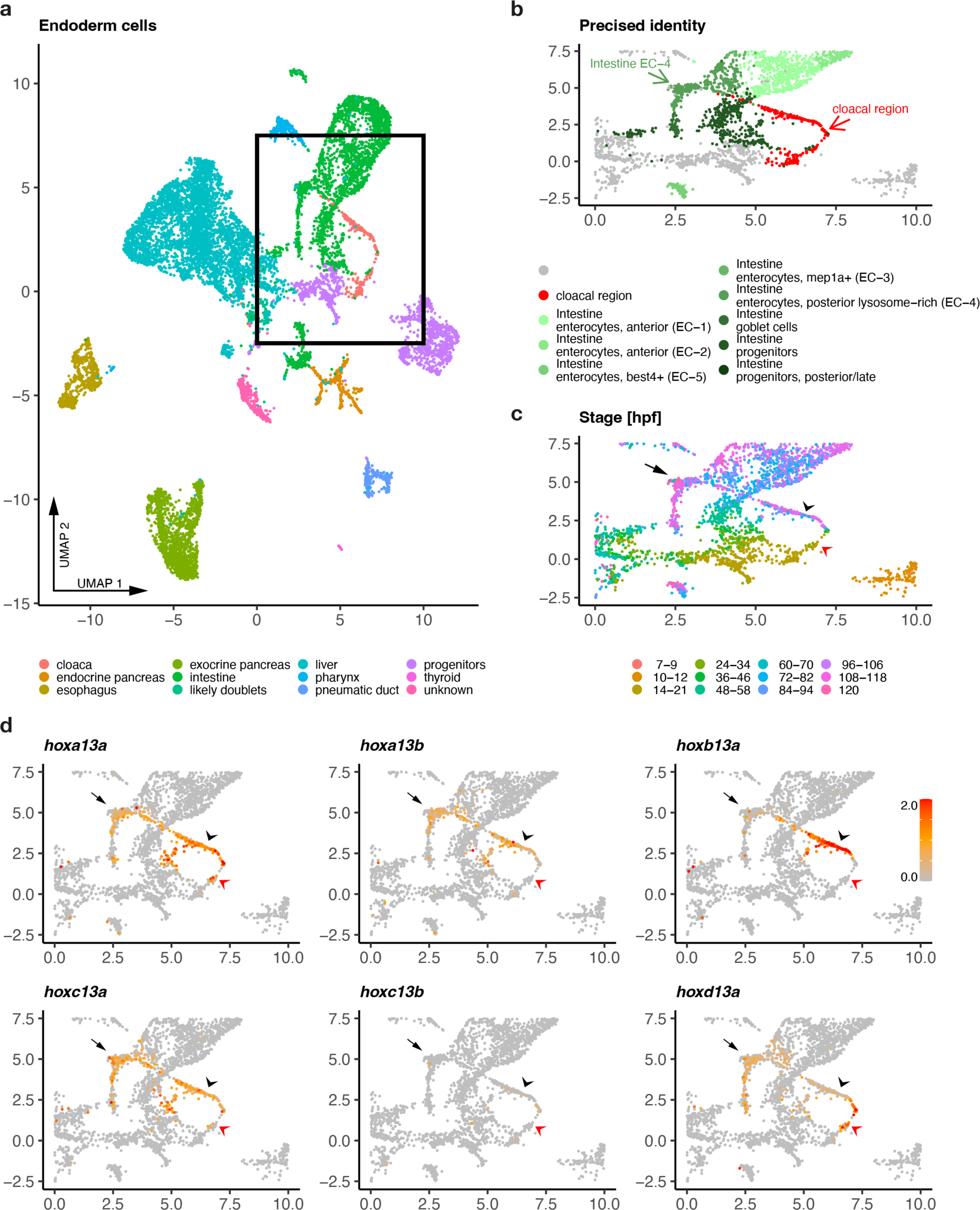
*hox13* gene expression in the Daniocell atlas. **a.** UMAP of endoderm cells using matrices extracted from ref.^56^, colored by tissue. The black rectangle indicates the UMAP region which contains cellular clusters from the cloacal region. All other panels in the figure correspond to this rectangle. **b.** UMAP of endodermal cells and identities of their clusters. The colors indicate the identities of cells from both the cloacal region (red arrow) and the posterior intestine (dark green, arrow). **c.** UMAP of selected endoderm, clustered by developmental stages (color code below). **d.** UMAP as in panel b, with the expression in red of the various *hox13* paralogous genes. All cells with a normalized expression level above 2 are displayed in red. In panels **c** and **d**, arrowheads indicate *hox13* expressing cells in the cloacal region either at early (red) or late (black) timepoints. The black arrows point to *hox13* expression in intestinal cells.

**Figure S11.**
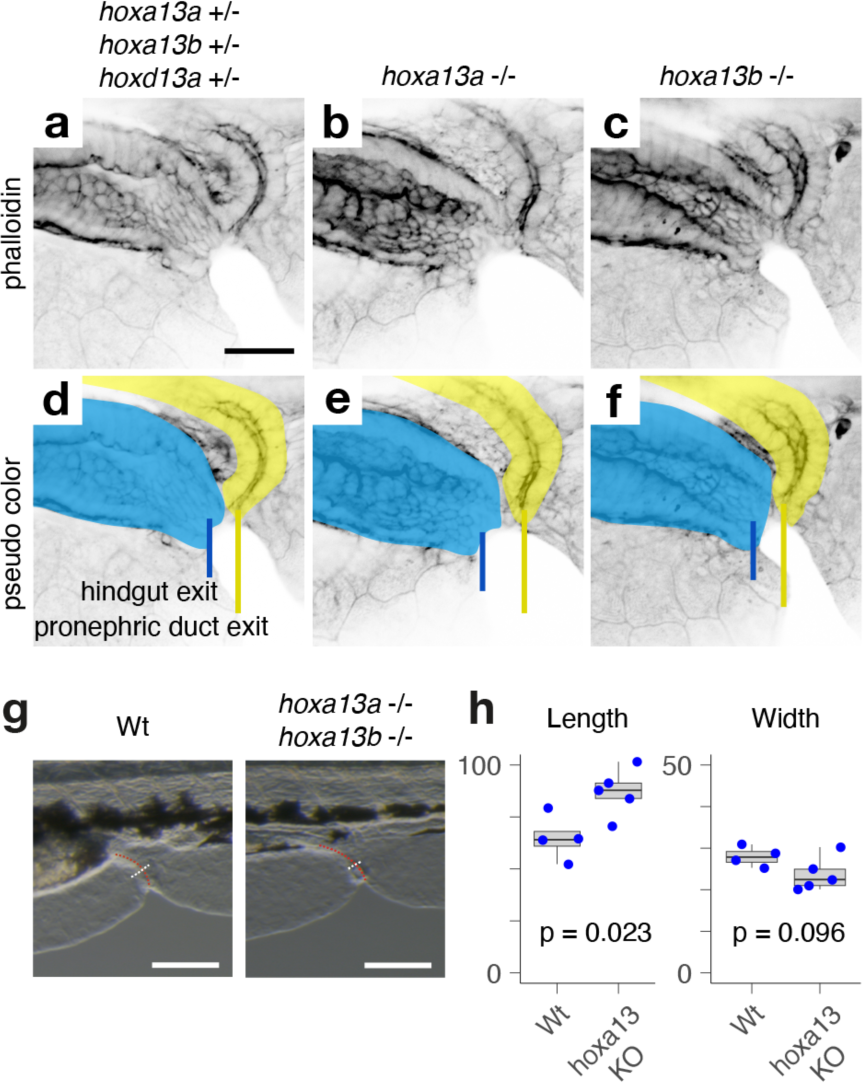
Cloacal region phenotypes in *hox13* mutant zebrafish. **a-f.** Confocal micrographs of mutant cloacal regions at 6 dpf shown in single channel (**a-c**) and pseudo color (**d-f**). **d.** Triple *hoxa13a*;*hoxa13b*;*hoxd13a* heterozygotes (n=6) exhibit wild-type patterning with separate openings for the hindgut (blue) and pronephric duct (yellow). **e.** Homozygous *hoxa13a* single mutants show wild-type patterning (n=4). **f.** Homozygous *hoxa13b* single mutants have wild-type patterning (n=4). **g-h.** Length and width of the hindgut and pronephric duct in wild-type and *hoxa13* mutant zebrafish embryos at 3 dpf. **g.** The length (red dotted lines) and width (white dotted lines) of the hindgut and pronephric complex at the median fin fold level were quantified in wild-type (n=4, left) and *hoxa13a*;*hoxa13b* double mutant embryos (n=5, right). **h.** The length difference of the hindgut and pronephric complex between wild-type and *hoxa13* double mutant embryos is statistically significant (\**p* = 0.0101, two-sided Welch’s t-test). The error bars indicate the standard error of the mean. Scale bar length is 30 μm in **a-f** and 100 μm in **g**.

**Table S1.** Extent of the mouse and zebrafish domains. Sizes are indicated in base pairs and were determined based on the transcription start sites of genes.

| mm39_short_name | mm39_RefSeq_ID | mm39_transcript_start_chr2 | danRer11_short_name | danRer11_RefSeq_ID | danRer11_transcript_start_chr9 |
| --- | --- | --- | --- | --- | --- |
| *Hoxd4* | NM_010469.2 | 74552322 | *hoxd4a* | NM_001126445.2 | 1951004 |
| *Hoxd13* | NM_008275.4 | 74498569 | *hoxd13a* | NM_131169.3 | 1990311 |
| *Nfe2l2* | NM_010902.5 | 75534860 | *nfe2l2a* | NM_182889.1 | 1654399 |
| *Atp5g3* | NM_001301721.1 | 73741670 | *atp5mc3a* | NM_201176.1 | 2333895 |

| Domain | Domain_name | mm39 | danRer11 | mm39/danRer11 |
| --- | --- | --- | --- | --- |
| *Atp5g3-Hoxd13* | 5DOM | 756899 | 343584 | 2.2 |
| *Hoxd4-Nfe2l2* | 3DOM | 982538 | 296605 | 3.3 |
| *Hoxd13-Hoxd4* | cluster | 53753 | 39307 | 1.4 |
| *whole genome* | assembly_Gb | 2.7 | 1.4 | 1.9 |

| Ratios | mm39 | danRer11 |
| --- | --- | --- |
| 5DOM/cluster | 14.1 | 8.7 |
| 3DOM/cluster | 18.3 | 7.5 |
| 5DOM/3DOM | 0.8 | 1.2 |

| Name | Assembly | Ratio |
| --- | --- | --- |
| 5DOM/cluster | mm39 | 14.1 |
| 3DOM/cluster | mm39 | 18.3 |
| 5DOM/3DOM | mm39 | 0.8 |
| 5DOM/cluster | danRer11 | 8.7 |
| 3DOM/cluster | danRer11 | 7.5 |
| 5DOM/3DOM | danRer11 | 1.2 |
| 5DOM | mm39/danRer11 | 2.2 |
| 3DOM | mm39/danRer11 | 3.3 |
| cluster | mm39/danRer11 | 1.4 |
| assembly | mm39/danRer11 | 1.9 |

| Assembly | Annotation source | Release |
| --- | --- | --- |
| danRer11 | NCBI RefSeq genes, curated subset | Annotation Release NCBI Danio rerio Annotation Release 106 (2019-10-28) |
| mm39 | NCBI RefSeq genes, curated subset (NM_*, NR_*, NP_* or YP_*) | Annotation Release NCBI RefSeq GCF_000001635.27-RS_2023_04 (2023-04-11) |

**Table S2.** Accession numbers for re-analyzed data. SRA accession numbers and reference of publications for the re-analyzed data when previously published data was used.

| Name | SRA number | Publication | DOI |
| --- | --- | --- | --- |
| CTCF_ChIP_E105_PT_rep1 | SRR17750150 | (Hintermann et al. 2022) | https://doi.org/10.1242/dev.200594 |
| Franke_CTCF_ChIP_24hpf_rep1 | SRR12435909 | (Franke et al. 2021) | 10.1038/s41467-021-25604-5 |
| Franke_CTCF_ChIP_24hpf_rep2 | SRR14670351 | (Franke et al. 2021) | 10.1038/s41467-021-25604-5 |
| Franke_CTCF_ChIP_48hpf_rep1 | SRR14670354 | (Franke et al. 2021) | 10.1038/s41467-021-25604-5 |
| Franke_CTCF_ChIP_48hpf_rep2 | SRR14670355 | (Franke et al. 2021) | 10.1038/s41467-021-25604-5 |
| Franke_48hpf_wt_rep1 | SRR12435867 | (Franke et al. 2021) | 10.1038/s41467-021-25604-5 |
| Franke_48hpf_wt_rep2 | SRR12435868 | (Franke et al. 2021) | 10.1038/s41467-021-25604-5 |
| Franke_24hpf_wt_rep1 | SRR14670388 | (Franke et al. 2021) | 10.1038/s41467-021-25604-5 |
| Franke_24hpf_wt_rep2 | SRR14670389 | (Franke et al. 2021) | 10.1038/s41467-021-25604-5 |
| Wike_24hpf_seq1 | SRR12044304 | (Wike et al. 2021) | 10.1101/gr.269860.120 |
| Wike_24hpf_seq2 | SRR12044305 | (Wike et al. 2021) | 10.1101/gr.269860.120 |
| Wike_24hpf_seq3 | SRR12044306 | (Wike et al. 2021) | 10.1101/gr.269860.120 |
| Wike_24hpf_seq4 | SRR12044307 | (Wike et al. 2021) | 10.1101/gr.269860.120 |
| Wike_24hpf_seq5 | SRR12044308 | (Wike et al. 2021) | 10.1101/gr.269860.120 |
| Wike_24hpf_seq6 | SRR12044309 | (Wike et al. 2021) | 10.1101/gr.269860.120 |

**Table S3.**
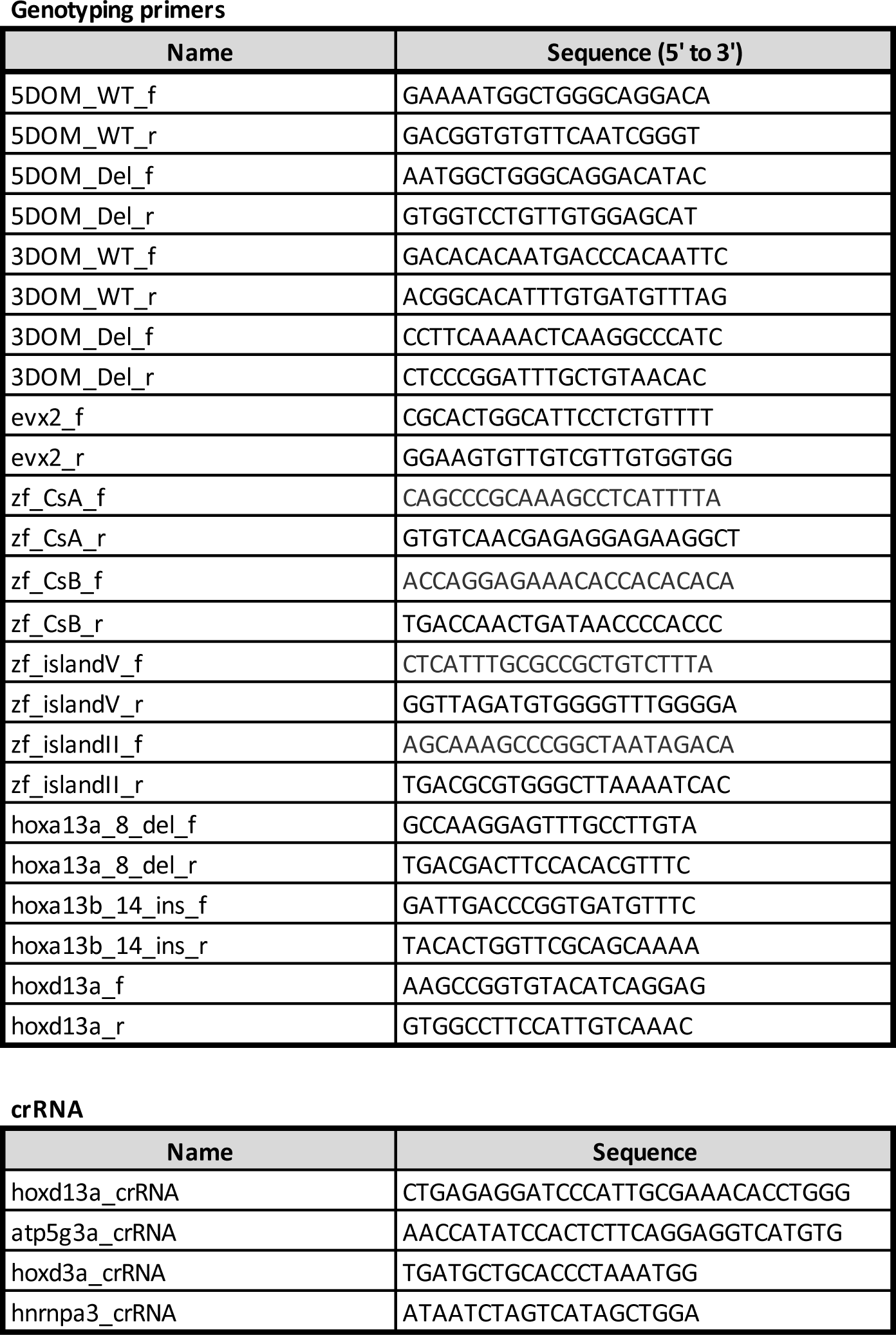
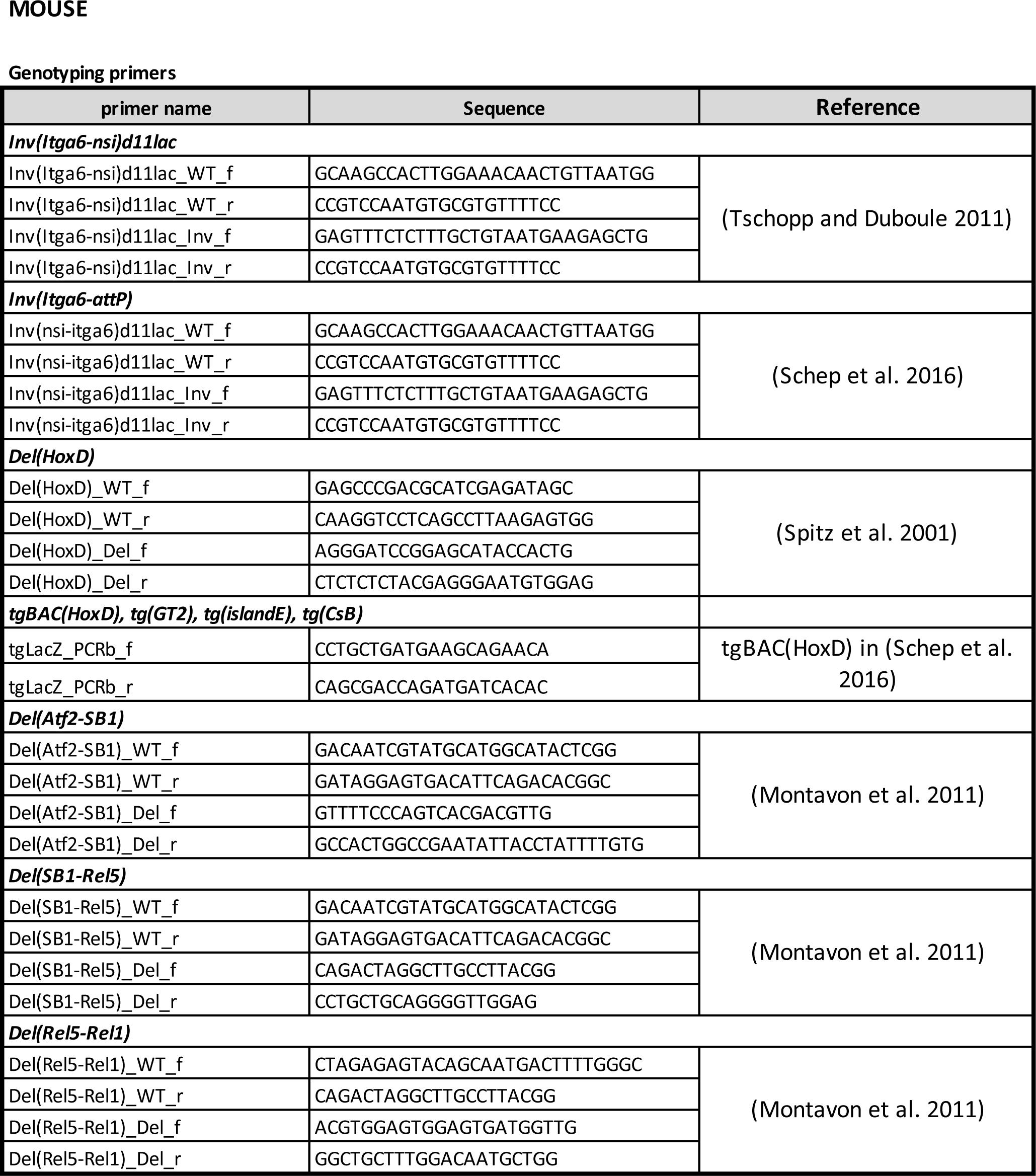
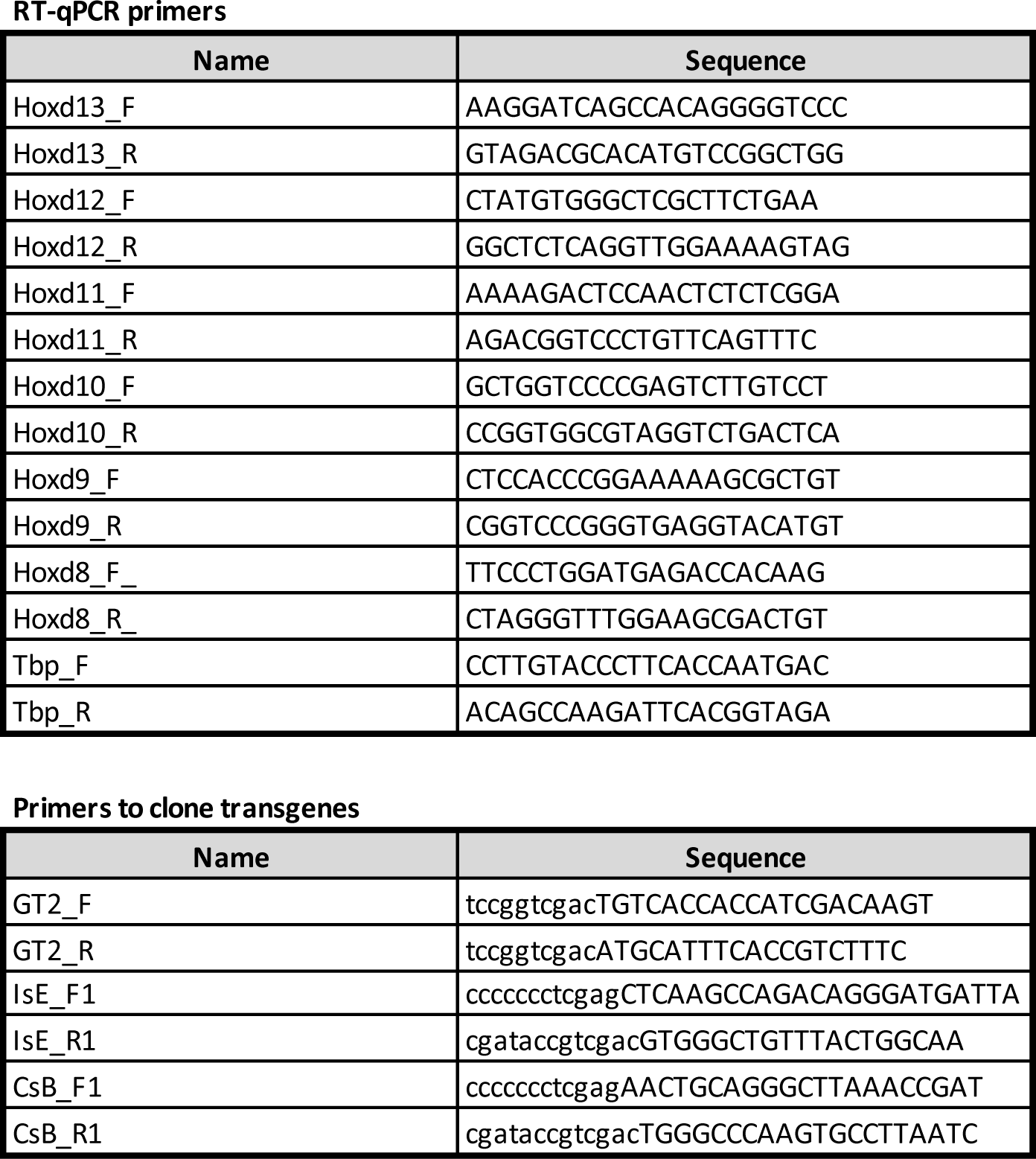
RT-qPCR primers. Lists of primers used in RT-qPCR experiments.

**File S1.** Sequences of the zebrafish probes used for WISH

**File S2.** Sequences of the mouse probes used for WISH

File S3. Sequences of the zebrafish *hoxda^Del^*(3DOM) and *hoxda^Del^*(5DOM) founder alleles

## MATERIALS AND METHODS

### Animal husbandry and ethics

All experiments using mice were approved and performed in compliance with the Swiss Law on Animal Protection (LPA) under license numbers GE45/20 and GE81/14. All animals were kept as a continuous backcross with C57BL6 × CBA F1 hybrids. Mice were housed at the University of Geneva Sciences III animal colony with light cycles between 07:00 and 19:00 in the summer and 06:00 and 18:00 in winter, with ambient temperatures maintained between 22 and 23 °C and 45 and 55% humidity, the air was renewed 17 times per hour. Zebrafish (*Danio rerio)* were maintained according to standard conditions^57^ under a 14/10 hours on/off light cycle at 26°C, with a set point of 7.5 and 600uS for pH and conductivity respectively. All zebrafish husbandry procedures have been approved and accredited by the federal food safety and veterinary office of the canton of Vaud (VD-H23). AB, Tu and TL were used as wild-type strains and were obtained from the European zebrafish resource center (EZRC). *hoxda^Del^*(3DOM) and *hoxda^Del^*(5DOM) mutants were generated for this study. Zebrafish embryos were derived from freely mating adults. Wild-type siblings, *hoxda^Del^*(3DOM) and *hoxda^De^*l(5DOM) homozygous embryos were obtained from crossing the corresponding heterozygous mutant. Embryos were collected within 30 minutes after spawning and incubated at 28.5°C in fish water, shifted at 20°C after reaching 80% epiboly and grown at 28.5°C to the proper developmental stage according to^58^. Pigmentation was prevented by treating the embryos with 0.002% N-phenylthiourea from 1 dpf onwards.

### Generation of the *hoxda^Del^*(3DOM) and *hoxda^Del^*(5DOM) deletions in zebrafish

The *hoxda^Del^*(3DOM) and *hoxda^Del^*(5DOM) mutant alleles were generated using the CRISPR/Cas9 system described in^59^. The sequences of the crRNAs used are listed in Table S3. Loci were identified with the GRCz11 zebrafish genome assembly available on Ensembl. The corresponding genomic regions were amplified and sequenced from fin clips. Adults carrying verified target sequences were isolated and then selected for breeding to generate eggs for genome editing experiments. The gRNAs target sites were determined using the open-source software CHOPCHOP (http://chopchop.cbu.uib.no/index.php) and chemically synthesized *Alt-R^®^* crRNAs and *Alt-R^®^* tracrRNAs as well as the *Alt-R^®^* Cas9 protein were obtained from Integrated DNA Technologies (IDT). To test the efficiency of these gRNAs in generating the expected mutant alleles, we injected boluses ranging from 100 µm to 150 µm and containing 5μM of the duplex crRNAs, tracrRNA and Cas9 RNP complex into the cytoplasm of one-cell stage embryos. Injecting the RNP complex solution in a 100 µm bolus gave less than 5% mortality. With this condition, 30% of the embryos carried the 5DOM deletion and 15% carried the 3DOM deletion. For each condition we extracted the genomic DNA of 20 individual larvae at 24hpf for genotyping^60^. Identification of *hoxda^Del^*(3DOM) and *hoxda^Del^*(5DOM) mutants were performed by PCR. Amplification of *evx2* was used as a control to confirm the presence or absence of the 5DOM. The PCR mix was prepared for using the Phusion® High-Fidelity DNA Polymerase (NEB) and primer sequences are listed in Table S3. In parallel, a hundred and twenty larvae per allele were raised to adulthood. To identify founders, F0 adults were outcrossed with wild-type and 25 embryos were genotyped. Three and four independent founders were obtained for the *hoxda^Del^*(5DOM) allele and *hoxda^Del^*(3DOM), respectively. Two founders of each deletion were verified by Sanger sequencing (File S3) and used for further experiments.

### Zebrafish *hox13* mutant lines, phalloidin labeling and genotyping

Frameshift loss-of-function alleles *hoxa13a^ch^*^307^, *hoxa13b^ch^*^308^, and *hoxd13a*^5b*p*^ *^ins^* were previously generated^9^. Zebrafish lines were propagated and maintained following^61^. To generate compound *hox13* mutants, animals triple heterozygous for *hoxa13a*, *hoxa13b* and *hoxd13a* were inter-crossed. Resulting larvae were fixed at 6 dpf in 4% paraformaldehyde in phosphate buffered saline (PBS) for 2 hours at room temperature with rocking agitation. After fixation, larvae were rinsed twice for 5 minutes each in PBS with added 1% Triton (PBSX). To visualize cloacal anatomy by labeling filamentous actin, larvae were then incubated in PBSX with fluorophore-conjugated phalloidin (Sigma-Aldrich P1951, phalloidin-Tetramethylrhodamine B isothiocyanate) added to a final concentration of 5U/mL overnight at 4°C with rocking agitation. Larvae were then rinsed twice for one hour each with PBSX.

For genotyping, phalloidin-labeled larvae were cut in half, separating the head, yolk, and pectoral fins from the cloaca and tail. The head half was used for genotyping and the tail half was saved at 4°C for later analysis. DNA was extracted from the head half by digesting tissue in proteinase K diluted to 1 mg/mL in 20 μl 1x PCR buffer (10 mM Tris-HCl, 50 mM KCl, 1.5 mM MgCl2) for 1 hour at 55°C followed by heat inactivation at 80°C for 20 minutes. The digested tissue was then subjected to brief vortexing and then 1 μl was used directly as template for genotyping PCR, with primers listed in Table S3. For thermocycling, after an initial step at 94°C for 2 minutes, reactions were cycled 40x though (15 s at 94°C, 15 s at 58 °C, 20 s at 72 °C) and finished with 5 minutes at 72 °C. PCR products were then heteroduplexed on a thermocycler by heating to 95°C for 10 minutes and then gradually cooled by 1°C every 10 seconds until a final temperature of 4°C was reached. Heteroduplexed PCR amplicons were then run on a high percentage agarose gel to determine genotype by product size.

To analyze cloacal morphology, fixed phalloidin-labeled tails were imaged on a Zeiss LSM 800 confocal microscope. After acquiring a full confocal stack through the cloacal region, a midline frame that demonstrated hindgut and pronephric duct morphology was selected.

### Mutant mouse stocks

All mouse lines used in this study were previously reported: *Inv(Itga6-nsi)d11lac*^33^, *Inv(Itga6-attP)* and *tgBAC(HoxD)*^32^, *Del(HoxD)*^34^ and *Del(Atf2-SB1)*, *Del(SB1-Rel5)* and *Del(Rel5-Rel1)*^62^.

### Whole-mount *in situ* hybridization (WISH)

The zebrafish and mouse antisense probes used in this study are listed in File S1 and File S2, respectively. For zebrafish, WISH were performed as described in^60^, at 58°C for all riboprobes (hybridization temperature and SSC washes). Wholemount embryos were photographed using a compound microscope (SZX10, Olympus) equipped with a Normarski optics and a digital camera (DP22, Olympus). Genotyping of individual embryos was performed after photographic documentation as in^60^ with primers listed in Table S3. Wild-type and mutant embryos originated from the same clutch of eggs produced by heterozygote crosses and underwent WISH procedure in the same well. Murine urogenital systems were isolated from E18.5 embryos and processed following a previously reported WISH procedure^63^, with some specific adjustments. For the Proteinase K treatment, urogenital systems were incubated 20 minutes in proteinase K diluted to 20 µg/ml in PBST. For the refixation step, a solution of 4% PFA containing 0.2% glutaraldehyde was used. Hybridization temperature was 69°C and temperature of post-hybridization washes was 65°C. Staining was developed in BM-Purple (Roche #11442074001) for approximately 4 hours at room temperature.

### Mouse genotyping

For extemporaneous genotyping, yolk sacs were collected and placed into 1.5 ml tubes containing Rapid Digestion Buffer (10 mM EDTA pH8.0 and 0.1 mM NaOH), then placed in a thermomixer at 95 °C for 10 min with shaking at 900 rpm. While the yolk sacs were incubating, the PCR master mix was prepared using Z-Taq (Takara R006B) and primers (Table S1) and aliquoted into PCR tubes. The tubes containing lysed yolk sacs were then placed on ice to cool briefly and quickly centrifuged at high speed. The lysate (1μl) was placed into the reaction tubes and cycled 32× (2 s at 98 °C, 2 s at 55 °C, 15 s at 72 °C). Twenty microliters of the PCR reaction were loaded onto a 1.5% agarose gel and electrophoresis was run at 120 V for 10 min. When samples could be kept for some time, a conventional genotyping protocol was applied with a Tail Digestion Buffer (10 mM Tris pH8.0, 25 mM EDTA pH8.0, 100 mM NaCl, 0.5% SDS) added to each yolk sac or tail clipping at 250μl along with 4μl Proteinase K at 20 mg/ml (EuroBio GEXPRK01-15) and incubated overnight at 55 °C. The samples were then incubated at 95 °C for 15 min to inactivate the Proteinase K and stored at −20 °C until ready for genotyping. Genotyping primers (Supplementary Data 1) were combined with Taq polymerase (Prospec ENZ-308) in 25μl reactions and cycled 2× with Ta = 64 °C and then cycled 32× with Ta = 62 °C.

### Mouse RT-qPCR

Urogenital sinuses (UGS) were collected from E18.5 male embryos separately and placed into 1× DEPC-PBS on ice. A little piece of the remaining embryo was collected for genotyping. The UGS were transferred into fresh 1× DEPC-PBS and then placed into RNALater (ThermoFisher AM7020) for storage at −80 °C until processing. Batches of samples were processed in parallel to collect RNA with Qiagen RNeasy extraction kits (Qiagen 74034). After isolating total RNA, first strand cDNA was produced with SuperScript III VILO (ThermoFischer 11754-050) using approximately 500 ng of total RNA input. cDNA was amplified with Promega GoTaq 2× SYBR Mix and quantified on a BioRad CFX96 Real Time System. Expression levels were determined by dCt (GOI–Tbp) and normalized to one for each condition by subtracting each dCT by the mean dCT for each wild-type set. Finally, expression was evaluated by two power this normalized dCT. Table S1 contains the primer sequences used for quantification. RT-qPCR measurements were taken from distinct embryos. Box plots for expression changes and two-tailed unequal variance t tests were produced in DataGraph 4.6.1. The boxes represent the interquartile range (IQR), with the lower and upper hinges denoting the first and third quartiles (25^th^ and 75^th^ percentiles). Whiskers extend from the hinges to the furthest data points within 1.5 times the IQR. The upper whisker reaches the largest value within this range, while the lower whisker extends to the smallest value within 1.5 times the IQR from the hinge.

### Mouse RNA-Seq

E18.5 male and female UGS were collected with a dissection separating the bladder from the urogenital sinus but including the proximal urethra (and vagina in females). Tissues were stored in RNALater (ThermFisher AM7020) and processed in parallel with Qiagen RNeasy extraction kits (Qiagen 74034). RNA quality was assessed on an Agilent Bioanalyzer 2100, with RIN scores > 9.5. RNA sequencing libraries were prepared at the University of Geneva genomics platform using Illumina TruSeq stranded total RNA with Ribo-Zero TM Gold Ribo-deleted RNA kits to produce strand-specific 100bp single-end reads on an Illumina HiSeq 2000. Raw RNA-seq reads were processed with CutAdapt version 4.1 (-a GATCGGAAGAGCACACGTCTGAACTCCAGTCAC -q 30 -m 15)^64^ to remove TruSeq adapters and bad quality bases. Filtered reads were mapped on the mouse genome mm39 with STAR version 2.7.10a^65^ with ENCODE parameters with a custom gtf file^66^ based on Ensembl version 108. This custom gtf file was obtained by removing readthrough transcripts and all noncoding transcripts from a protein-coding gene. FPKM values were evaluated by Cufflinks version 2.2.1^67,68^ with options --max-bundle-length 10000000 --multiread-correct --library-type “fr-firststrand” -b mm10.fa --no-effective-length-correction -M MTmouse.gtf -G. Boxplots depicting expression levels in distinct embryos were generated using the same methodology as for RT-qPCR.

### ATAC-Seq

Mouse and fish tissues were isolated and placed into 1x PBS containing 10% FCS on ice. Collagenase (Sigma-Aldrich C9697) was added to 50ug/ml and incubated at 37° for 20 minutes with shaking at 900rpm. Cells were washed 3x in 1x PBS. The number of cells was counted and viability confirmed to be greater than 90%. An input of 50000 cells was then processed according to previous description^21^. Sequencing was performed at EPFL GECF on an Illumina NextSeq 500. Analysis was performed similarly to^69^. Raw ATAC-seq paired-end reads were processed with CutAdapt version 4.1 (-a CTGTCTCTTATACACATCTCCGAGCCCACGAGAC -A CTGTCTCTTATACACATCTGACGCTGCCGACGA -q 30 -m 15)^64^ to remove Nextera adapters and bad quality bases. Filtered reads were mapped on mm39 for mouse samples and danRer11 where alternative contigs were removed for fish samples with bowtie2 version 2.4.5^70^ with the following parameters: --very-sensitive --no-unal --no-mixed --no-discordant --dovetail -X 1000. Only pairs mapping concordantly outside of mitochondria were kept (Samtools v1.16.1)^71^. PCR duplicates were removed by Picard version 3.0.0 (http://broadinstitute.github.io/picard/index.html). BAM files were converted to BED with bedtools version 2.30.0^72^. Peaks were called and coverage was generated by MACS2 version 2.2.7.1 with --nomodel --keep-dup all --shift -100 --extsize 200 --call-summits -B. Coverages were normalized to million mapped reads.

### Mouse ChIP-Seq

Male UGS were isolated and placed into 1x PBS containing 10% FCS on ice. ChIP-seq experiments were then performed as previously described in^73^. Briefly, they were fixed for 10 mn in 1% formaldehyde at room temperature and the crosslinking reaction was quenched with glycine. Subsequently, nuclei were extracted and chromatin was sheared using a water-bath sonicator (Covaris E220 evolution ultra-sonicator). Immunoprecipitation was done using the following anti-H3K27ac (Abcam, ab4729) or anti-H3K27me3 (Merck Millipore, 07–449). Libraries were prepared using the TruSeq protocol, and sequenced on the Illumina HiSeq 4000 (100 bp single-end reads) according to manufactures instructions. CTCF re-analysis from^69,74^. Accession numbers are listed in TableS2. Raw ChIP-seq single-reads or paired-end reads were processed with CutAdapt version 4.1 (-a GATCGGAAGAGCACACGTCTGAACTCCAGTCAC for single-reads and -a CTGTCTCTTATACACATCTCCGAGCCCACGAGAC -A CTGTCTCTTATACACATCTGACGCTGCCGACGA -q 30 -m 15)^64^ to remove Truseq or Nextera adapters and bad quality bases. Filtered reads were mapped on mm39 for mouse samples and danRer11 where alternative contigs were removed for reanalysis of fish samples with bowtie2 version 2.4.5^70^ with the default parameters. Only alignments with a mapping quality above 30 were kept (Samtools v1.16.1)^75^. Peaks were called and coverage was generated by MACS2 version 2.2.7.1 with with --call-summits -B (and --nomodel --extsize 200 for single-read). Coverages were normalized to million mapped reads/pairs.

### Mouse enhancer-reporter assay

Transgenic embryos were generated as described in^35^. Primers were designed to amplify genomic DNA from the region around the observed ATAC and H3K27Ac peaks (Table S3). These primers included extra restriction sites for either *XhoI* or *SalI* at the 5’ ends. The PCR fragments were cleaned with Qiagen Gel Extraction Kit (#28704). The PCR fragment and the pSKlacZ reporter construct (GenBank X52326.1)^73^ were digested with *XhoI* or *SalI* and ligated together with Promega 2× Rapid Ligation kit (#C6711). Sanger sequencing confirmed the correct sequences were inserted upstream of the promoter. Maxipreps of the plasmid were prepared and eluted in 1x IDTE (#11-05-01-13). Pro-nuclear injections were performed and embryos were collected at approximately E18.5 and stained for *lacZ*. UGS were collected from E18.5 embryos in ice-cold 1× PBS in a 12-well plate. All steps were done with gentle shaking on a rocker plate at room temperature. Tissues were fixed for 5 min at room temperature in freshly prepared 4% PFA. After fixing, they were washed three times in 2 mM MgCl2, 0.01% Sodium Deoxycholate, 0.02% Nonidet P40, and 1× PBS, for 20 min at RT. The wash solution was replaced by X-gal staining solution (5 mM Potassium Ferricynide, 5 mM Potassium Ferrocynide, 2 mM MgCl2 hexahydrate, 0.01% Sodium Deoxycholate, 0.02% Nonidet P40, 1 mg/ ml X-Gal, and 1× PBS) for overnight incubation with the plate wrapped in aluminum foil to protect from light. Tissues were then washed three times in 1× PBS and fixed in 4% PFA for long-term storage. Images of embryos were collected with an Olympus DP74 camera mounted on an Olympus MVX10 microscope using the Olympus cellSens Standard 2.1 software.

### Mouse Capture HiC-seq

E18.5 male UGS were collected and collagenase-treated samples were cross-linked with 1% formaldehyde (Thermo Fisher 28908) for 10 min at RT and stored at −80° until further processing as previously described^76^. The SureSelectXT RNA probe design used for capturing DNA was done using the SureDesign online tool by Agilent. Probes cover the region chr2: 72240000–76840000 (mm9) producing 2X coverage, with moderately stringent masking and balanced boosting. DNA fragments were sequenced on the Illumina HiSeq 4000 and processed with HiCUP version 0.9.2 on mm39 with --re1 GATC^77^, bowtie2 version 2.4.5^70^ and samtools version 1.16.1^75^. The output BAM was converted to pre-juicer medium format with hic2juicer from HiCUP. The pairs with both mates on chr2:72233000–76832000 were selected, sorted and loaded into a 10kb bins matrix with cooler version 0.8.11^78^. The final matrix was balanced with the option --cis-only. The TADs were computed with HiCExplorer hicFindTADs version 3.7.2^79,80^ with --correctForMultipleTesting fdr --minDepth 120000 --maxDepth 240000 --step 240000 --minBoundaryDistance 250000. Data was plotted on mm39 (chr2:73600000-75550000).

### Zebrafish HiC-seq

The HiC profiles derived from a reanalysis of data from^55,74^. Accession numbers are listed in Table S2. Reads were mapped on danRer11 where alternative contigs were removed and no selection of reads were performed. Valid pairs were loaded into a 10kb bins matrix. TAD calling parameters were adapted to the smaller size of the genome: --chromosomes “chr9” --correctForMultipleTesting fdr --minDepth 35000 --maxDepth 70000 --step 70000 --minBoundaryDistance 50000. Data was plotted on danRer11 (chr9:1650000-2400000) and on an inverted *x* axis.

## CUT&RUN

Zebrafish samples were processed according to (10.1038/nprot.2018.015), using a final concentration of 0.02% digitonin (Apollo APOBID3301). Approximately 0.5e6 cells were incubated with 0.1μg/100μl of anti-H3K27ac antibody (Abcam Ab4729), or 0.5 μg/100 μl of anti-H3K27me3 (Merck Millipore 07-449) in Digitonin Wash Buffer at 4 °C. The pA-MNase was kindly provided by the Henikoff lab (Batch #6) and added at 0.5 μl/100 μl in Digitonin Wash Buffer. Cells were digested in Low Calcium Buffer and released for 30 min at 37 °C. Sequencing libraries were prepared with KAPA HyperPrep reagents (07962347001) with 2.5ul of adapters at 0.3uM and ligated for 1 h at 20 °C. The DNA was amplified for 14 cycles. Post-amplified DNA was cleaned and size selected using 1:1 ratio of DNA:Ampure SPRI beads (A63881) followed by an additional 1:1 wash and size selection with HXB. HXB is equal parts 40% PEG8000 (Fisher FIBBP233) and 5 M NaCl. Sequencing was performed at EPFL GECF on an Illumina HiSeq 4000. Raw CUT&RUN paired-end reads were processed with CutAdapt version 4.1 (-a GATCGGAAGAGCACACGTCTGAACTCCAGTCAC -A GATCGGAAGAGCGTCGTGTAGGGAAAGAGTGT -q 30 -m 15) to remove TruSeq adapters and bad quality bases^64^. Filtered reads were mapped on danRer11 where alternative contigs were removed with bowtie2 version 2.4.5^70^ with the following parameters: --very-sensitive --no-unal --no-mixed --no-discordant --dovetail -X 1000. Only alignments with mapping quality above 30 were kept (Samtools v1.16.1)^75^. PCR duplicates were removed by Picard version 3.0.0 (http://broadinstitute.github.io/picard/index.html). BAM files were converted to BED with bedtools version 2.30.0^72^. Peaks were called and coverage was generated by MACS2 version 2.2.7.1 with -- nomodel --keep-dup all --shift -100 --extsize 200 --call-summits -B. Coverages were normalized to million mapped reads.

### Analyses of conserved sequences

Annotation of orthologous domains was performed using transcription start sites of orthologous genes, as reported in Table S1. To identify conserved sequences between mouse and zebrafish, a pairwise alignment was done between the mouse genomic region chr2:73600000-75550000 (mm39) and the zebrafish orthologous region chr9:1650000-2400000 (danRer11) using discontinuous mega blast. To reduce false positives, only reciprocal hits were considered. To display multispecies conservation levels, MAF files were generated between the chromosome 2 of the mouse genome (mm39) and contig chrUn_DS181389v1 of the Platypus genome (ornAna2), chromosome 7 of the chicken genome (galGal6), contig chrUn_GL343356 of Lizard genome (anoCar2), chromosome 9 of the frog genome (xenTro10), contig JH127184 of the Coelacanth genome (latCha1), chromosome 9 of the zebrafish genome (danRer11), chromosome 1 of the Fugu genome (fr3) and the whole lamprey genome (petMar3). Details for the maf generation are available on the github repository https://github.com/AurelieHintermann/HintermannBoltEtAl2024. To help the visualization, a horizontal line was plotted for each species on each region.

### scRNA-seq

The matrix of the scRNA-seq atlas was downloaded from GEO (GSE223922)^56^ as well as the table with metadata. The matrix was loaded into a Seurat object with Seurat version 4 3.0^81^ with R version 4.3.0. Cells attributed to the ‘tissue.name’ ‘endoderm’ were selected. Normalization and PCA was done as in ref.^56^ and UMAP was performed on the top 70 PCA and 50 nearest neighbors. UMAP coordinates and *hox13* normalized expression of endoderm cells were exported to file and plotted with ggplot2 version 3.4.4.

### Software

The phylogenic tree was generated with http://timetree.org using the following species: *Mus musculus, Protopterus, Danio rerio, Carcharhinus leucas, Petromyzon marinus, Branchiostoma lanceolatum* and subsequently edited with seaview 4.7. Genomic tracks from Next-Generation Sequencing were plotted with pyGenomeTracks 3.8, using custom gene annotations available at https://zenodo.org/records/7510797 (mm39) and https://zenodo.org/records/10283274 (danRer11). RT-qPCR, RNA-seq and domain size quantifications were plotted with R using the ggplot package.

## ACKNOWLEDGEMENTS

We thank all members of the Duboule laboratories for comments and discussions, the Van der Goot laboratory for providing a fish cDNA library, Mikiko Tanaka for the *hoxd10a* probe, Jeffrey Farrell for his advices and Andy Oates for his support. The calculations were performed using the facilities of the Scientific IT and Application Support Center of EPFL. This work was supported in part using the resources and services of the Gene Expression Research Core Facility (GECF) at the School of Life Sciences of EPFL and transgene injections were performed at the transgenesis core facility of the University of Geneva, medical school. We acknowledge the support of the Brinson Family Foundation (to NS).

## AUTHORS CONTRIBUTIONS

Conceptualization: AH, CCB, DD

Methodology: AH, CCB, GV, LLD, BM, MBH

Investigation: AH, CCB, GV, LLD, SG, PBG, BM, TN, TAM, MBH

Visualization: AH, CCB, GV, DD, MBH, TN, TAM

Funding acquisition: DD, MPH, NS

Project administration and supervision: AH, CCB, DD

Writing the original draft: AH, CCB, DD

Review writing & editing: AH, DD, MPH, MBH, NS

## ETHICAL STATEMENT

All experiments involving animals were performed in agreement with the Swiss Law on Animal Protection (LPA). For mice, work was carried out under license No GE 81/14 (to DD). For zebrafish, work was carried out under a general license of the EPFL granted by the Service de la Consommation et des Affaires Vétérinaires of the canton of Vaud, Switzerland (No VD-H23).

## DATA AVAILABILITY

All raw and processed datasets are available in the Gene Expression Omnibus (GEO) repository under accession number GSE250267.

## CODE AVAILABILITY

All scripts necessary to reproduce figures from raw data are available at https://github.com/AurelieHintermann/HintermannBoltEtAl2024.

## Sequences for zebrafish ISH probes

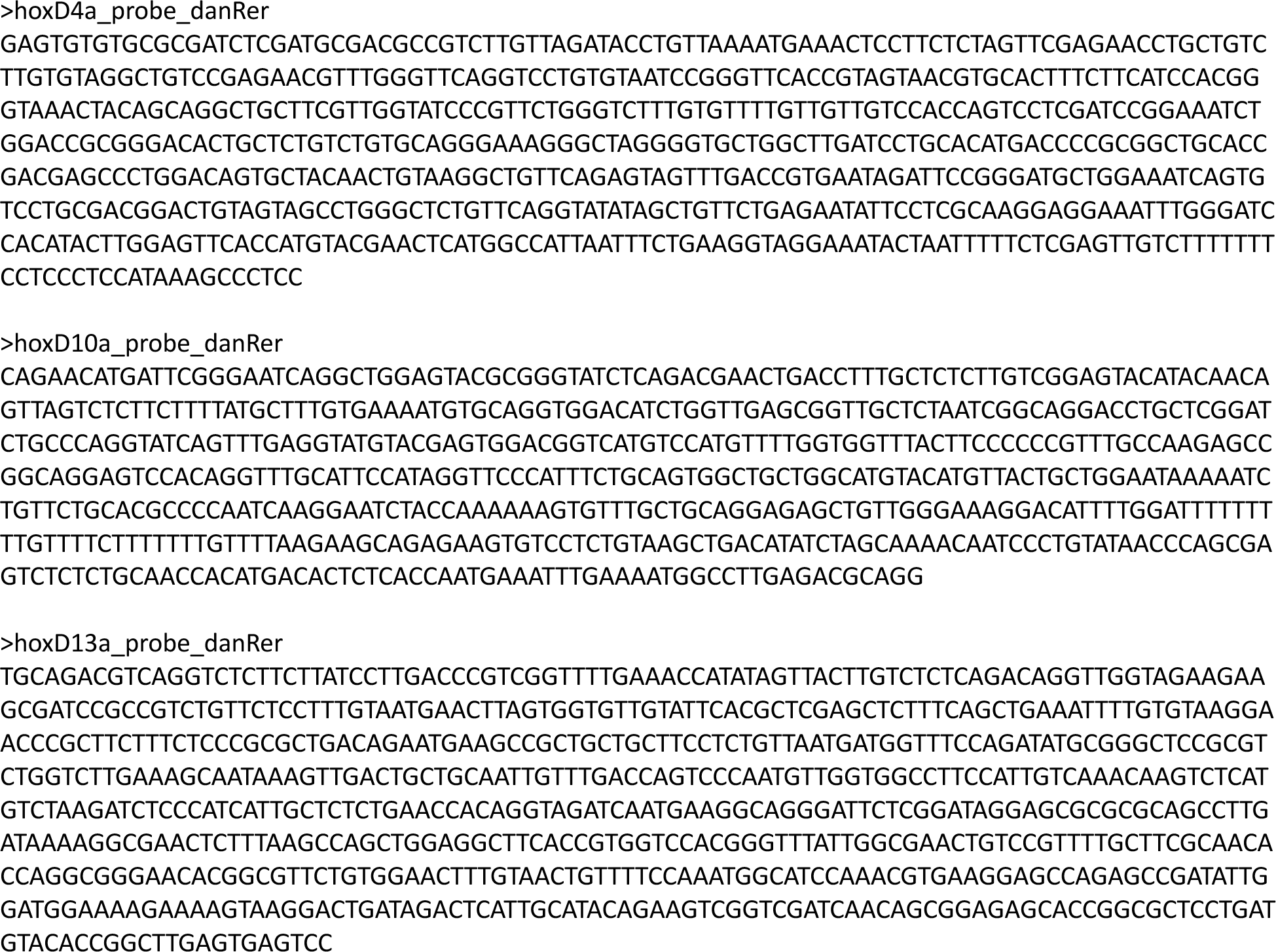

## Sequences for mouse ISH probes

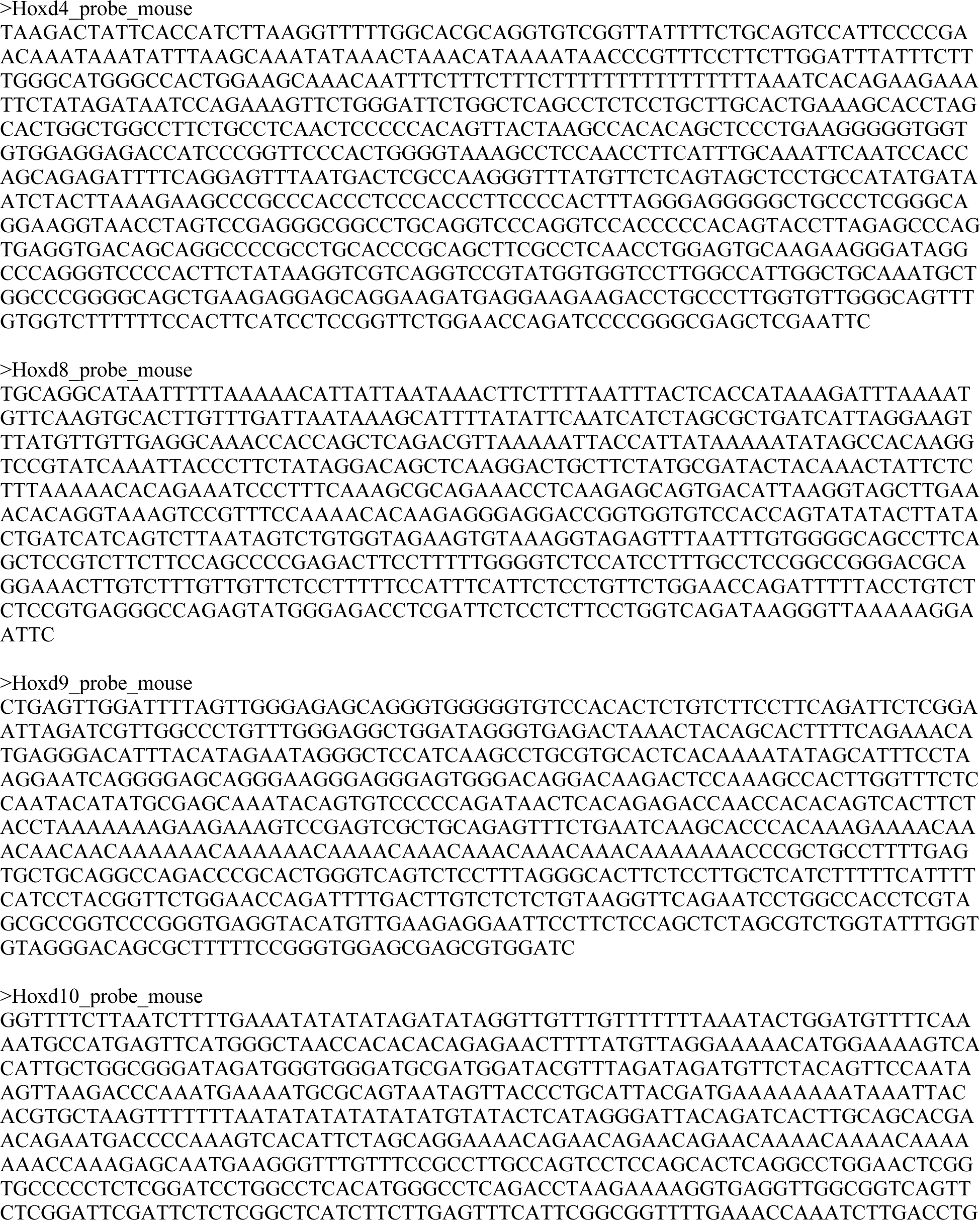

## Sanger sequences of the zebrafish founders for *hoxda^Del^*(3DOM) and *hoxda^Del^*(5DOM)

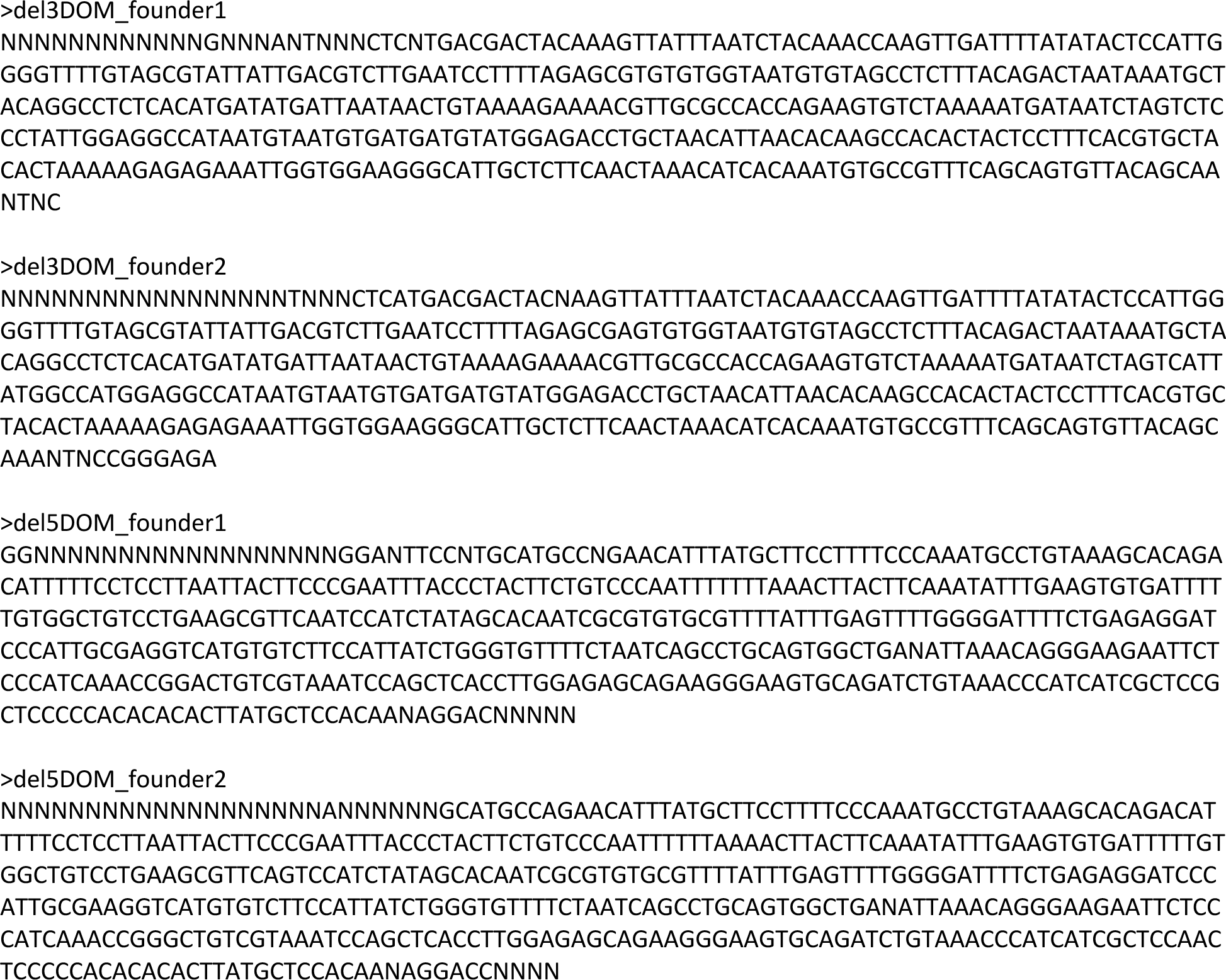

